# Cameroonian blackflies (Diptera: Simuliidae) harbour a plethora of (RNA) viruses

**DOI:** 10.1101/2024.04.05.588247

**Authors:** Lander De Coninck, Amber Hadermann, Ludovica Ingletto, Robert Colebunders, Kongnyu Gamnsi Njamnshi, Alfred Kongnyu Njamnshi, John L Mokili, Joseph Nelson Siewe Fodjo, Jelle Matthijnssens

## Abstract

Strong epidemiological evidence suggests that onchocerciasis may be associated with epilepsy – hence the name onchocerciasis-associated epilepsy (OAE). However, the pathogenesis of OAE still needs to be elucidated, as recent studies failed to detect *Onchocerca volvulus* in the central nervous system of persons with OAE. Therefore, it was suggested that a potentially neurotropic virus transmitted by blackflies could play a role in triggering OAE. To investigate this hypothesis, adult blackflies were collected in an onchocerciasis-endemic area with a high OAE prevalence in the Ntui Health District, Cameroon. A whole-genome shotgun sequencing approach was used to detect viral sequences in fifty-five pools of ten blackflies. A very high abundance of viral reads was detected across multiple (novel) viral families, including viral families associated with human disease. Although, no genomes closely related to known neurotropic viruses were found in the blackfly virome, the plethora of novel viruses representing novel species, genera and even families, warrant further exploration for their potential to infect vertebrates. These results could serve as a first step for studying the viruses associated with the hematophagous blackfly, which also could be present in their nematode host *O. volvulus*. Exploring the diversity of viruses in blackflies should be included in the active surveillance of zoonotic diseases.

## Introduction

Human onchocerciasis, commonly known as river blindness, is a parasitic disease caused by the filarial worm *Onchocerca volvulus* (Nematoda: Secernentea; Spirurida; Filariidae; Onchocerca). Transmission of the parasite occurs through repeated bites of female blackflies belonging to the *Simulium* species (Brattig, Cheke and Garms 2021). Onchocerciasis is classified as one of the neglected tropical diseases recognized by the World Health Organization (WHO).

Onchocerciasis-endemic regions with high ongoing or past *O. volvulus* transmission are known to have high epilepsy prevalence (Pion and Boussinesq 2012; Levick *et al*. 2017; Colebunders *et al*. 2018b; Siewe Fodjo *et al*. 2018; Boullé *et al*. 2019; Mukendi *et al*. 2019; Raimon *et al*. 2021). While onchocerciasis is commonly thought to affect only the skin and eyes, emerging epidemiological evidence suggests a potential direct or indirect association between *O. volvulus* infection and epilepsy, giving rise to the term “Onchocerciasis-Associated Epilepsy” (OAE), also known as “River Epilepsy” (Chesnais *et al*. 2018, 2020; Colebunders *et al*. 2019, 2021; Hadermann *et al*. 2023). OAE presents with a wide spectrum of epileptic seizures, including generalized tonic-clonic seizures, myoclonic seizures, absence seizures and head nodding seizures (Colebunders *et al*. 2018a). Nodding and Nakalanga syndrome are considered to be phenotypic presentations of OAE (Colebunders *et al*. 2018a; Van Cutsem *et al*. 2023) Nakalanga syndrome is characterized by severe growth retardation, delayed sexual development, mental impairment, facial deformation, kyphoscoliosis, and epileptic seizures (Föger *et al*. 2017; Siewe Fodjo *et al*. 2019).

So far, the pathogenesis of OAE is not yet established. Prior to the introduction of ivermectin mass drug distribution, O. volvulus microfilariae (mf) had been detected in the cerebrospinal fluid (CSF) of individuals with onchocerciasis (Duke, Vincelette and Moore 1976). However, in more recent studies, neither O. volvulus mf nor DNA could be detected in CSF (Winkler *et al*. 2013; Hotterbeekx *et al*. 2020) nor in post-mortem brain tissue of persons with OAE (Hotterbeekx *et al*. 2019). Therefore, it was suggested that a potentially neurotropic virus transmitted by blackflies, or an endosymbiont of the *O. volvulus* parasite could play a role in the pathogenesis of OAE ((Colebunders *et al*. 2014; Hadermann *et al*. 2023).

Very little is known about the virome of blackflies. Kraberger *et al*. described the ssDNA virome of blackflies in New Zealand and reported high numbers of *Genomoviridae*, *Circoviridae* and *Microviridae* (Kraberger *et al*. 2019). Moreover, invertebrate iridoviruses (family *Iridoviridae*), have been identified in blackfly larvae across the globe (Piégu *et al*. 2013, 2014a, 2014b). In addition, the most studied arbovirus transmitted by blackflies is vesicular stomatitis virus (family *Rhabdoviridae;* genus *Vesiculovirus*), that typically infects cattle, horses, and swine (Howerth, Mead and Stallknecht 2002). Nonetheless, there have been no reports of blackflies transmitting arboviruses to humans to date. However, considering the 1700 *Simulium* species described worldwide, it is plausible that blackflies may transmit unidentified viruses from human and zoonotic origins. The aim of this study was to identify potential neurotropic viruses (in this study defined as viruses related to viruses known to cause infections in the central nervous system, like West Nile virus, Japanese encephalitis virus [*Flaviviridae*] and rabies virus [*Rhabdoviridae*]) transmitted by blackflies, which could play a role in the pathogenesis of OAE.

## Materials and Methods

### Sample collection

Adult blackflies investigated were collected in the village of Nachtigal (coordinates of the breeding site: N 4°21.146, E 11°37.953; Garmin GPSMAP78). The gathering took place from 7:00 AM to 6:00 PM over three consecutive days in July 2021. Nachtigal is a rural community of the Ntui Health District, an onchocerciasis-endemic area with a high prevalence of OAE (Siewe Fodjo *et al*. 2022), in the forest-savannah transition zone in the Centre Region of Cameroon. This community lives close to the Nachtigal rapids on the Sanaga River, which constitutes one of the main sources of blackfly vectors in Cameroon (Siewe Fodjo *et al*. 2022). The human landing catch method was used to collect the blackflies and mouth aspirators were used to aspirate blackflies into an empty container every hour (Hendy *et al*. 2021). At the end of each catching day, blackflies were stored dry at −20°C in the BRAIN laboratory, Yaoundé, Cameroon until shipping on dry ice a few weeks later. The frozen blackfly samples were then sent to the Laboratory of Viral Metagenomics in Leuven, Belgium, where they were stored at −80°C until the nucleic acids were extracted for sequencing.

### Sample processing

A total of fifty-five pools of ten blackflies were analysed. The NetoVIR protocol (Conceição-Neto *et al*. 2015) was adapted for the homogenization of the blackfly pools. Homogenization was carried out in 2mL tubes with 2.8mm zirconium oxide beads at 5000 rpm for 15 seconds. Negative controls were added in the beginning and taken through the whole process (including sequencing) to make sure no contamination happened during the processing of the samples. Samples were centrifuged and filtered through a 0.8 µm filter (Sartorius) to remove cellular debris and enrich the samples for viral particles. This filtrate was treated with a mix of Benzonase (50 U, Novagen) and Micrococcal nuclease (2000 U, New England Biolabs) to digest free-floating nucleic acids. DNA and RNA was simultaneously extracted using the QiaAMP Viral RNA Mini kit (Qiagen) without carrier RNA. A random amplification of both DNA and RNA with the Whole Transcriptome Amplification (WTA2) kit increased the nucleic acid concentration of the samples, and resulting PCR products were further purified and prepared for sequencing with the Nextera XT kit (Illumina). The final sequencing libraries were cleaned up with Agencourt AMPure XP beads (Beckman Coulter, Inc.) using a 0.6 ratio of beads to sample. Paired-end sequencing (2×150bp) was performed on the Novaseq 6000 SP platform at the Nucleomics Core sequencing facility (VIB Leuven).

### Raw Reads Processing and Virome Analysis

To process the raw paired-end Illumina reads, an in-house bioinformatics pipeline (ViPER v1.1) (De Coninck 2021) was used with the ‘triple assembly’ option enabled. Briefly, adapters and low-quality reads were trimmed with Trimmomatic (Bolger, Lohse and Usadel 2014), before a *de novo* assembly with metaSPAdes v3.15.3 (Nurk *et al*. 2017). Resulting fasta files with scaffolds were subsequently concatenated. To remove redundancy in the data, these scaffolds were clustered on 95% nucleotide identity over 85% coverage of the shortest sequence with the clustering algorithm available from CheckV (Nayfach *et al*. 2020). A length filter of 1000 nucleotides was applied on the sequences before viruses were identified in the data using a combined approach. This approach consisted of: 1) genomad v1.7.0 on default setting with score calibration enabled (Camargo *et al*. 2023); 2) DIAMOND blastx v2.0.11 on the sensitive setting (Buchfink, Xie and Huson 2014) with NCBI’s nr database (accessed 23-03-2023); and 3) a Hidden Markov Model (HMM) search of the scaffolds’ open reading frames (ORFs), predicted by prodigal (Hyatt *et al*. 2010), against the NeoRdRP HMM dataset (v1.0) (Sakaguchi *et al*. 2022) with HHMER v3.3.2 (Eddy 2011) with a minimum e-value 1e-10. Taxonomy assignment for blastx-identified sequences was decided based on a lowest common ancestor approach from the best twenty-five hits. When possible, the genomad taxonomy output was prioritized for a sequence over the blastx taxonomic assignment. Subsequently, for quality control purposes, CheckV v1.0.1 (Nayfach *et al*. 2020) was employed to assess the genome completeness of the identified viral sequences. Sequences not belonging to the *Riboviria* and less than 50% complete as predicted by CheckV were removed from further analysis. *Riboviria* sequences predicted to be less than 50% complete were kept because CheckV does not handle (segmented) RNA viruses well, which often results in wrong completeness estimations for RNA viruses. Finally, using the sequencing data of the negative controls, sequences that were predicted by the prevalence method of decontam v1.22.0 (Davis *et al*. 2018) to be a contaminant were removed from our dataset.

### Phylogenetic Analysis

Based on the taxonomic assignment of the scaffolds (*vide supra*), we downloaded the International Committee on Taxonomy of Viruses (ICTV) representative protein sequences of each family or order present in our dataset (from ICTV’s MSL39 v1 VMR) with NCBI Virus. Except for the *Orthomyxoviridae* of which representative PB1 sequences were downloaded from https://github.com/evogytis/orthomyxo-metagenomics/blob/main/data/Fig3/PB1_full.fasta (Dudas and Batson 2023). For the RNA viruses, we first filtered out all non-RdRP proteins and partial RdRPs with a combination of palm_annot (https://github.com/rcedgar/palm_annot; commit 15d9443) and palmscan2 (https://github.com/rcedgar/palmscan) to retrieve the intact palmcore domain (Babaian and Edgar 2022). For the *Genomoviridae* and *Parvoviridae*, we downloaded the Rep and NS1 protein sequences, respectively. For each family or order, a diversified ensemble was created with Muscle5 (Edgar 2022). This diversified ensemble constitutes 100 multiple sequence alignments (MSAs) of the sequences with perturbed alignment parameters and permuted guide trees (see Muscle5 documentation for more information). From these 100 MSAs, the alignment with the highest column confidence, calculated with muscle maxcc, was picked to generate the final phylogenetic tree. Resampling of the diversified ensemble to generate support values for the phylogenetic tree was obtained with muscle resample - conf 0 -gapfract 0.5. Maximum likelihood (ML) phylogenetic trees from the resampled MSAs were inferred with Fasttree (Price, Dehal and Arkin 2010) (-lg -gamma -nosupport). These ML trees served as support for the tree that was generated from the maxcc MSA (also generated with Fasttree and unaltered settings), which was calculated with newick conftree (https://github.com/rcedgar/newick).

### Viral host inference

To infer if the *Flaviviridae* and *Rhabdoviridae* viruses in our data could have an invertebrate or vertebrate host, we calculated the relative dinucleotide abundance (RDA) of our sequences and sequences with a known host (see Supplementary Table S2 for accession numbers) with DinuQ (Lytras and Hughes 2020). Next, we performed a principal component analysis in R on all dinucleotide RDA values and plotted the first against the second principal component for all viruses.

### Ethical approval

Approval was obtained from the ethics committee of the Cameroon Baptist Convention Health Services (reference number IRB2021-03). We also obtained a research permit from the Ministry of Scientific Research and Innovation (Ref: 000144/MINRESI/B00/C00/C10/C13), as well as an ABS-Nagoya Protocol Prior Informed Consent (Ref: Decision No. 00016/D/MINEPDED/CNA) and ABS Permit No. 00013/MINEPDED/CAN/NP-ABS/ABS-FP) from the Ministry of Environment, Protection of Nature and Sustainable Development.

## Results

### Identification of more than thousand viral contigs

For the fifty-five blackfly pools sequenced, between 6,113,870 and 24,002,232 reads were obtained per pool. In total, around 720 million raw reads were generated from all fifty-five pools. Raw read classification with Kraken2 (Wood, Lu and Langmead 2019) showed that most reads could not be directly classified, followed by on average 30% of the reads assigned to Bacteria (See Supplemental Figure 1B). After trimming, each pool retained between 4,544,534 and 20,790,370 reads, resulting in a 17.5% loss of the initial raw reads (see Supplementary Table S1 and Supplemental Figure 1A). Subsequently, *de novo* assemblies were generated, resulting in a total of 59,645 scaffolds larger than 1000 bp from the 55 pools. The clustering of the assembled scaffolds from all samples based on 95% nucleotide identity over 85% coverage resulted in 45,308 non-redundant scaffolds. After removing contamination predicted with decontam, a total of 1,678 scaffolds were identified as viral with a combination of genomad, DIAMOND blastx and an HMM search against the NeoRdRP dataset (see Supplemental Figure 2A). Of these, 56 were considered to be 100% complete, 128 were of high quality (>90% complete) and 190 of medium quality (>50% complete), the remaining viral sequences were either of low quality or their completeness could not be determined by CheckV (see Supplemental Figure 2B). However, the low-quality and not-determined groups consisted only of sequences with *Riboviria* or unclassified taxonomic realm assignments. This suggests that these sequences are significantly divergent and might thus be of a higher quality than what is estimated by CheckV (also see Materials). Overall, we concluded that, due to the high convergence of our three identification methods and the completeness estimation of CheckV, we have a robust set of identified viruses.

### The blackfly virome is dominated by RNA viruses

When examining the taxonomic distribution of our viral genome sequences, the majority (n=1186) were classified within the *Riboviria* realm, 322 could not be assigned into a specific realm and a small number belonged to the *Monodnavaria* and the *Duplodnaviria* (see Figure 1A). At the read level, a distinct picture arises with approximately two thirds being identified as belonging to the *Duplodnaviria* realm. This is mostly because the average viral genome length of *Duplodnaviria* is much higher than the average genome length for the *Riboviria*, which in our data recruit about one third of the viral reads (see Figure 1A and Supplemental Figure 3). The 322 unclassified sequences recruit only a small fraction (∼1.5 million) of the reads. As the diversity seemed the greatest in the *Riboviria* realm, which also contains the vast majority of arboviruses, we further investigated the taxonomic distribution of their scaffolds as well as their reads. Around half of the viral genome sequences belonged to the *Duplornaviricota* phylum (see Figure 1B), while this phylum constituted only one third of the *Riboviria* reads (see Figure 1B). Surprisingly, 81 further unclassified *Riboviria* sequences were responsible for another third of the viral reads in our blackfly samples. These results show that the blackflies contained a high number of completely novel RNA viruses that were present in high abundance. Moreover, we constructed rarefaction curves to assess the depth of sampling required to encompass the full spectrum of viral diversity (Figure 1C). Notably, the curves (total and *Riboviria*) plateaued after approximately fifty pools, suggesting saturation and the absence of novel viral taxa beyond this point. These findings indicate that our sampling efforts comprehensively cover the viral diversity within the blackflies from the onchocerciasis-endemic Nachtigal village (Cameroon).

**Figure 1:**
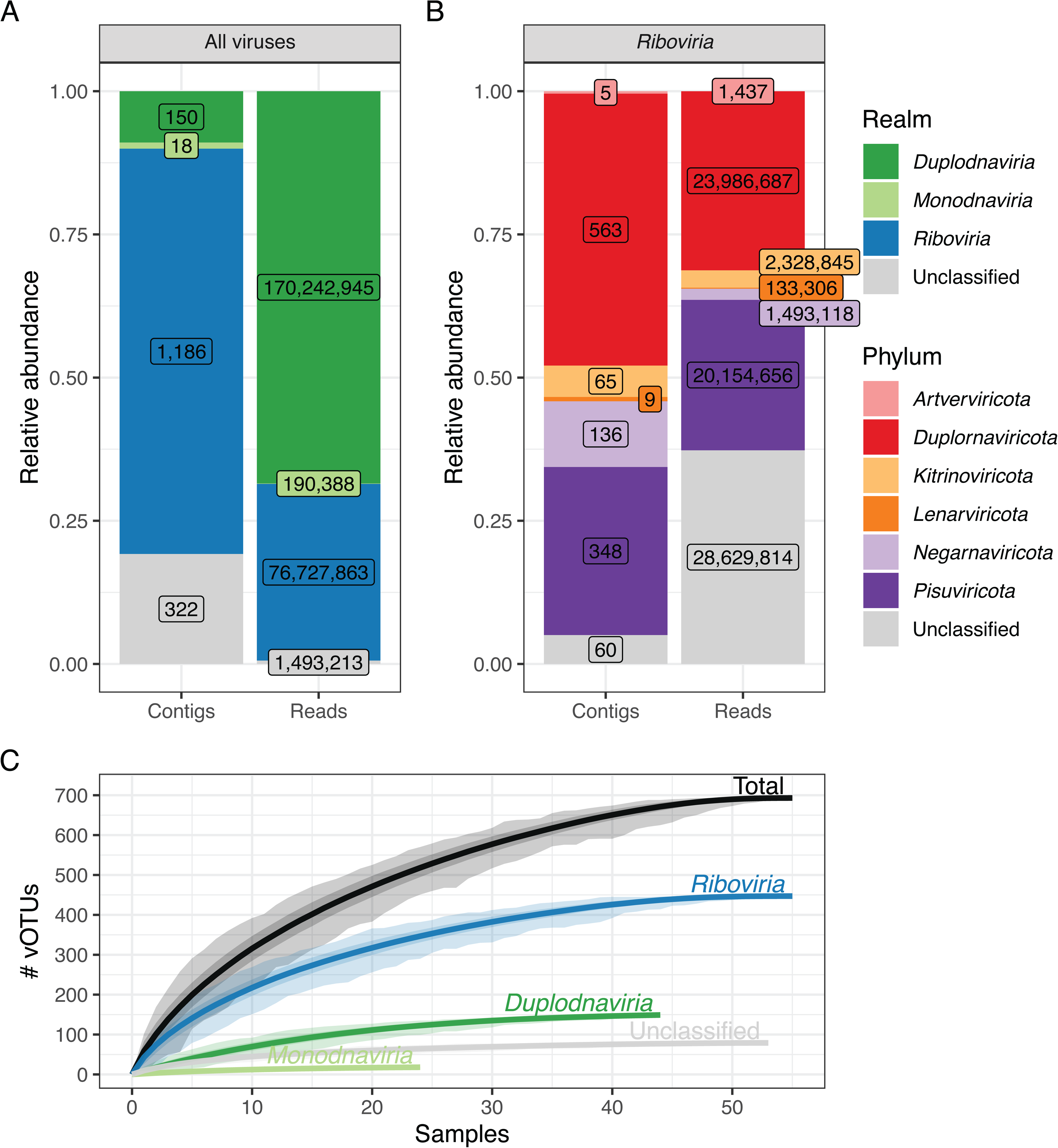
Blackfly viromes are dominated by RNA viruses. A) Barplots showing the relative abundance of all identified viruses at contig and read level. Labels indicate the absolute numbers. B) Barplots showing the relative abundance of all identified *Riboviria* phyla at contig and read level. Labels indicate the absolute numbers. C) Rarefaction curves for each virus kingdom and the total. Light shaded areas indicate the minimum and maximum observed species across all iterations, the dark shaded areas indicate the 95% confidence around the mean.

To visualize the virome similarity of the fifty-five pools, a heatmap with the read count for each viral family on a log₂ scale was constructed (see Figure 2). In total, forty eukaryotic viral families were identified across all samples. In addition, eleven different taxonomic instances of prokaryotic viruses were identified on the family level. Most diversity in the blackflies on the family level is present in the *Riboviria* realm with thirty-five families of eukaryotic RNA viruses detected.

**Figure 2:**
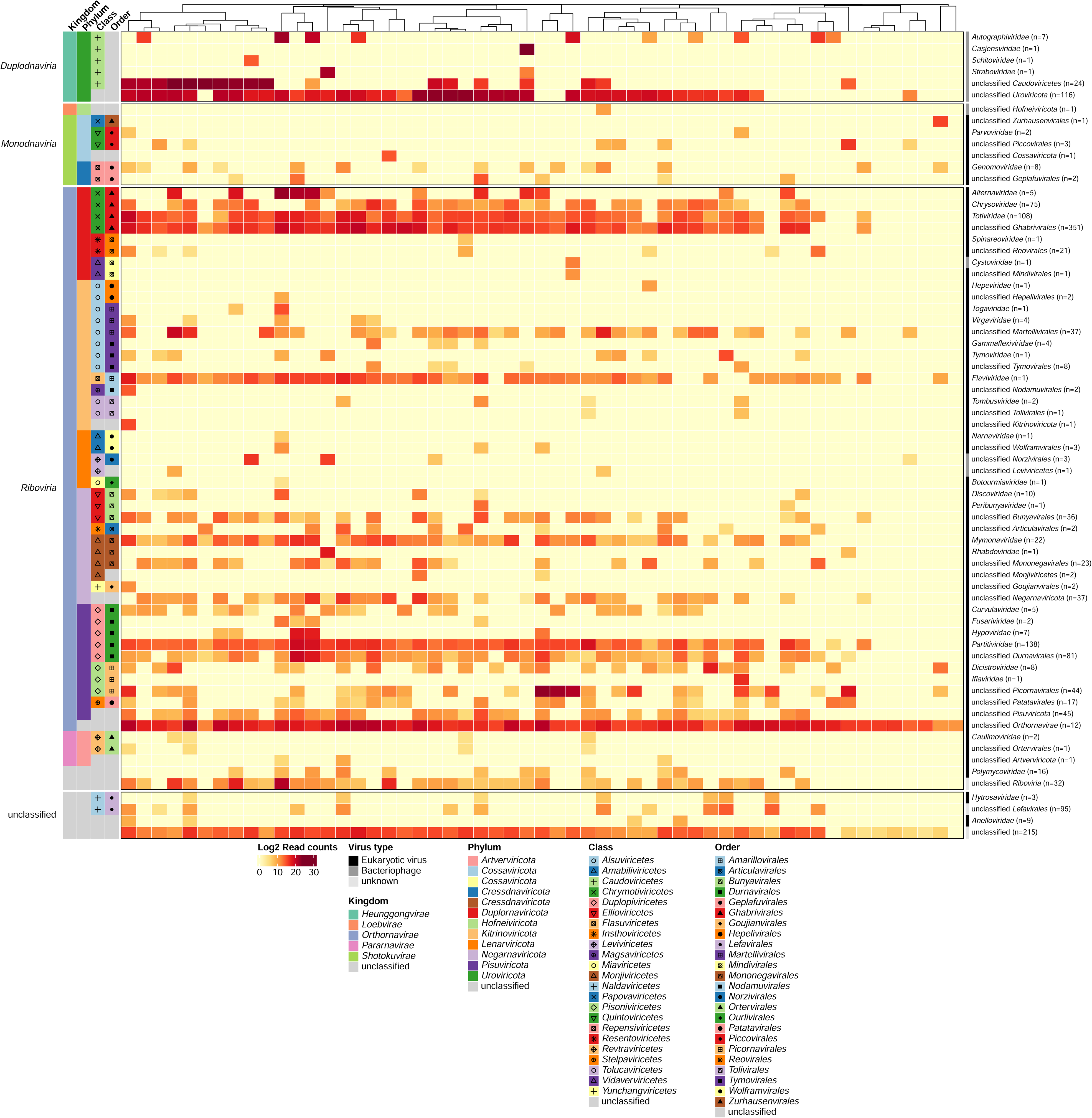
Blackflies harbour a large number of viral families, including families of potential clinical interest. Heatmap of all viral families per blackfly pool based on the blastx results (sequence similarity cutoff of at least 70% was used to assign scaffolds to families, otherwise they were assigned to closest related order). The heatmap shows the read count on a log2 scale for eukaryotic virus on family level from all the 55 adult blackfly pools. The number of identified scaffolds is indicated after each viral family name in the heatmap. Hierarchical clustering of the columns is based on the Bray-Curtis distance calculated from the read counts.

The viral families *Flaviviridae, Hepeviridae, Orthomyxoviridae, Parvoviridae, Phenuiviridae, Spinareoviridae, Rhabdoviridae*, and *Togaviridae* include human pathogenic viruses, some of which may also be arboviruses. Notably, known viruses causing neurological complications belong to the families of *Rhabdoviridae*, *Flaviviridae, Picornaviridae* as well as *Togaviridae*.

Fungal viruses were the most prevalent viruses, with members of *Ghabrivirales* (i.e., *Chrysoviridae*, *Totiviridae*) and *Partitiviridae* being the most abundant. A virus of the family *Flaviviridae* was present in most blackfly pools and in a relatively high abundance. Furthermore, other eukaryotic viral families (main hosts according to ICTV are given in brackets) that were prevalent, were *Curvulaviridae* (fungi), *Dicistroviridae* (invertebrates), *Iflaviridae* (insects), *Genomoviridae* (human, mammals, birds, fungi), *Mymonaviridae* (fungi), *Orthomyxoviridae* (human, birds, pigs), *Parvoviridae* (vertebrates, insects), *Rhabdoviridae* (vertebrates, invertebrates, plants), *Totiviridae* (fungi) and *Virgaviridae* (plants). From these families we constructed phylogenetic trees, either grouped in their respective order or as a separate family.

### RNA virus phylogenetics

To fully display the diversity of RNA viruses in the blackflies, we made phylogenetic trees of the palmcore sequences of the RdRp protein for the most common and most interesting RNA viruses present. These palmcore sequences might not completely reflect the true phylogeny of the viruses in the tree, but they prove to be a robust way to taxonomically classify sequences as shown by the high monophylicity of established families and genera (see Supplemental Figures 4-13).

The majority of the viruses in the blackflies seem fungi-infecting as they are part of the *Ghabrivirales*, *Durnavirales* (dsRNA) and *Mymonaviridae* (ssRNA-) and cluster together with ICTV exemplar viruses that infect fungi (see Figure 3 and Supplemental Figures 4, 5 and 6, respectively). Within the dsRNA orders, the blackfly virus sequences often form deep branches in the tree suggestive the discovery of new species, genera or maybe even families, thereby expanding the known diversity of the *Partitiviridae* and *Curvulaviridae* in the *Durnavirales*, and the *Chrysoviridae* and *Pseudototiviridae* within the *Ghabrivirales*.

**Figure 3:**
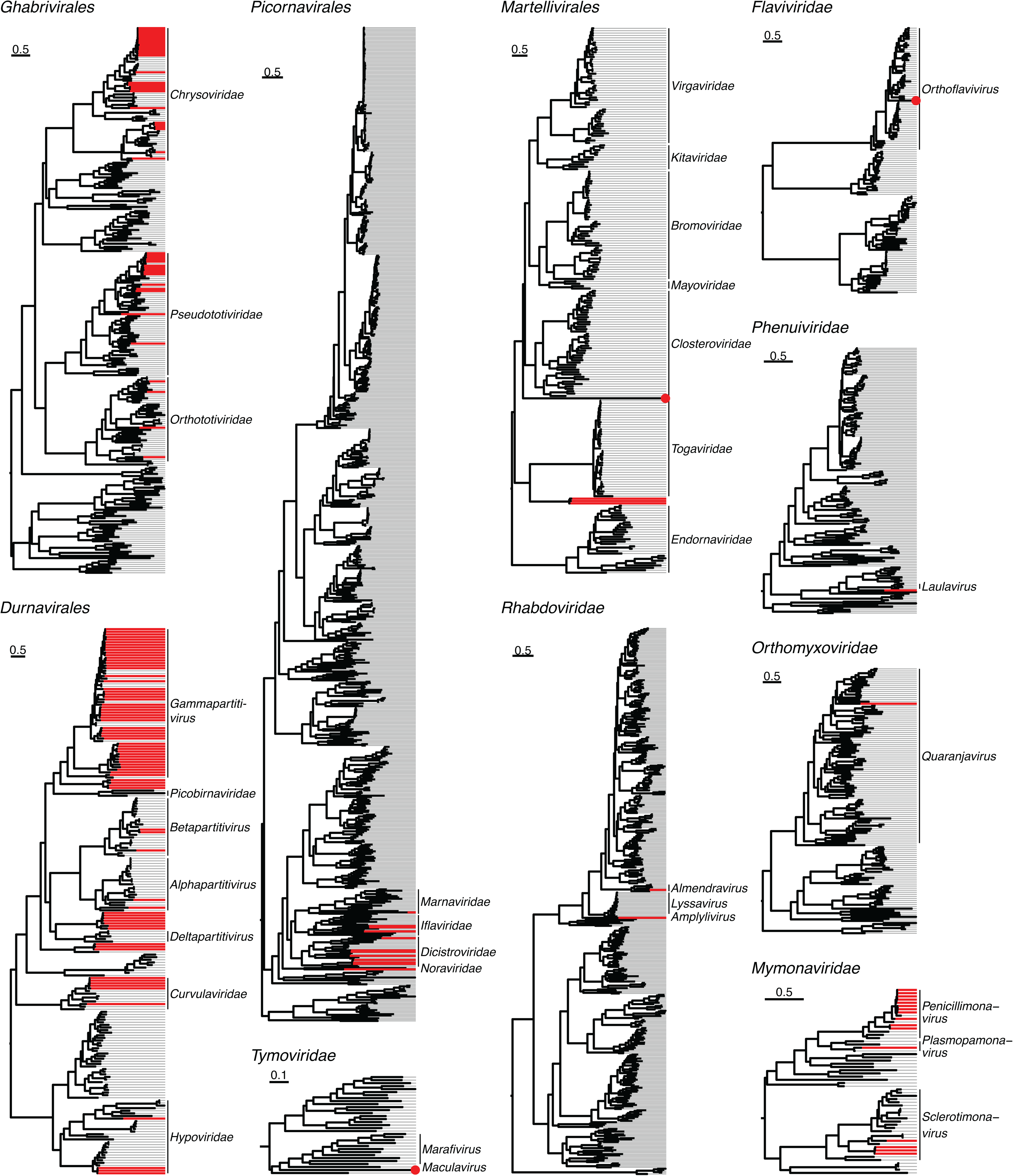
Phylogenetic trees representing the major RNA virus orders and families identified in the blackflies. Red lines signify novel viruses identified in the blackflies, while grey lines are ICTV references. The scale bars indicate the number of amino acid substitutions per site. Families and/or genera closest to the novel viruses are shown at the tips of the tree with a bar covering all ICTV exemplar species within that taxonomy (or no bar when there is only one exemplar species).

Viruses truly infecting the blackflies are mostly found in the *Picornavirales* order (ssRNA+), within the *Iflaviridae* and *Dicistroviridae* (see Figure 3 and Supplemental Figure 7). Furthermore, within the *Martellivirales*, which contains mostly plant-infecting viruses apart from the *Togaviridae*, we have possibly two new families. Interestingly, the viruses most closely related to the *Togaviridae* are probably segmented viruses as the recovered sequences only contain an RdRP gene. Attempts to retrieve more segments with a contig co-occurrence analysis (Batson *et al*. 2021) were unfortunately unsuccessful due to the low prevalence of these viruses in the blackfly samples. Also blastx searches against NCBI’s nr database revealed only sequences with only an RdRP gene and without matching segment in NCBI’s nt database coding for a structural protein/proteins.

Furthermore, we have a few singular viruses in the families *Tymoviridae*, *Flaviviridae*, *Phenuiviridae*, and *Orthomyxoviridae* of which the host is most likely the blackfly. In the *Rhabdoviridae*, we find two viruses: one that is clearly a member of the *Almendravirus* genus which contains invertebrate-infecting viruses and one virus that is not a part of a known genus but is part of a large clade that contains viruses infecting vertebrates and invertebrates as well as both (see Supplemental Figure 13).

### Novel blackfly rhabdo- and flaviviruses are most likely invertebrate-specific

To investigate whether the *Rhabdoviridae* and *Flaviviridae* member viruses we found in the blackflies could also infect vertebrates, we compared the dinucleotide composition within their genomes with the genomes of viruses with a known host. After coloring the viruses by host, this revealed a clear clustering in vertebrate-infecting vs. insect-specific viruses for the *Flaviviridae* (see Figure 4A), meaning that this family of viruses adapted itself to avoid the vertebrate immune system by using similar dinucleotide abundances in their genome (Lytras and Hughes 2020; Fros *et al*. 2021). For the *Rhabdoviridae* this clustering is less clear, but there seems to be nevertheless a small difference. The flavivirus we found in the blackflies shows a dinucleotide profile suggestive of being an insect-specific virus and also the rhabdoviruses we found in the blackflies seem to be likely infecting invertebrates when combining both the RDA information and the phylogenetic analysis (see Figure 4B and Supplemental Figure 13). Additionally, we added two other virus species recently identified in other studies to the *Rhabdoviridae* dinucleotide analysis: *Onchocerca volvulus* RNA virus 1 (OVRV1) is a rhabdovirus that was found in *Onchocerca volvulus* and elicits an immune response in the host (human as well as cattle and hamsters) (Quek *et al*. 2024), while Mundri virus was detected in a patient with nodding syndrome (Edridge *et al*. 2022). Interestingly, both multiple OVRV1 variants as well as Mundri virus seem to cluster with vertebrate infecting rhabdoviruses, suggesting that these viruses may be capable of infecting vertebrates.

**Figure 4:**
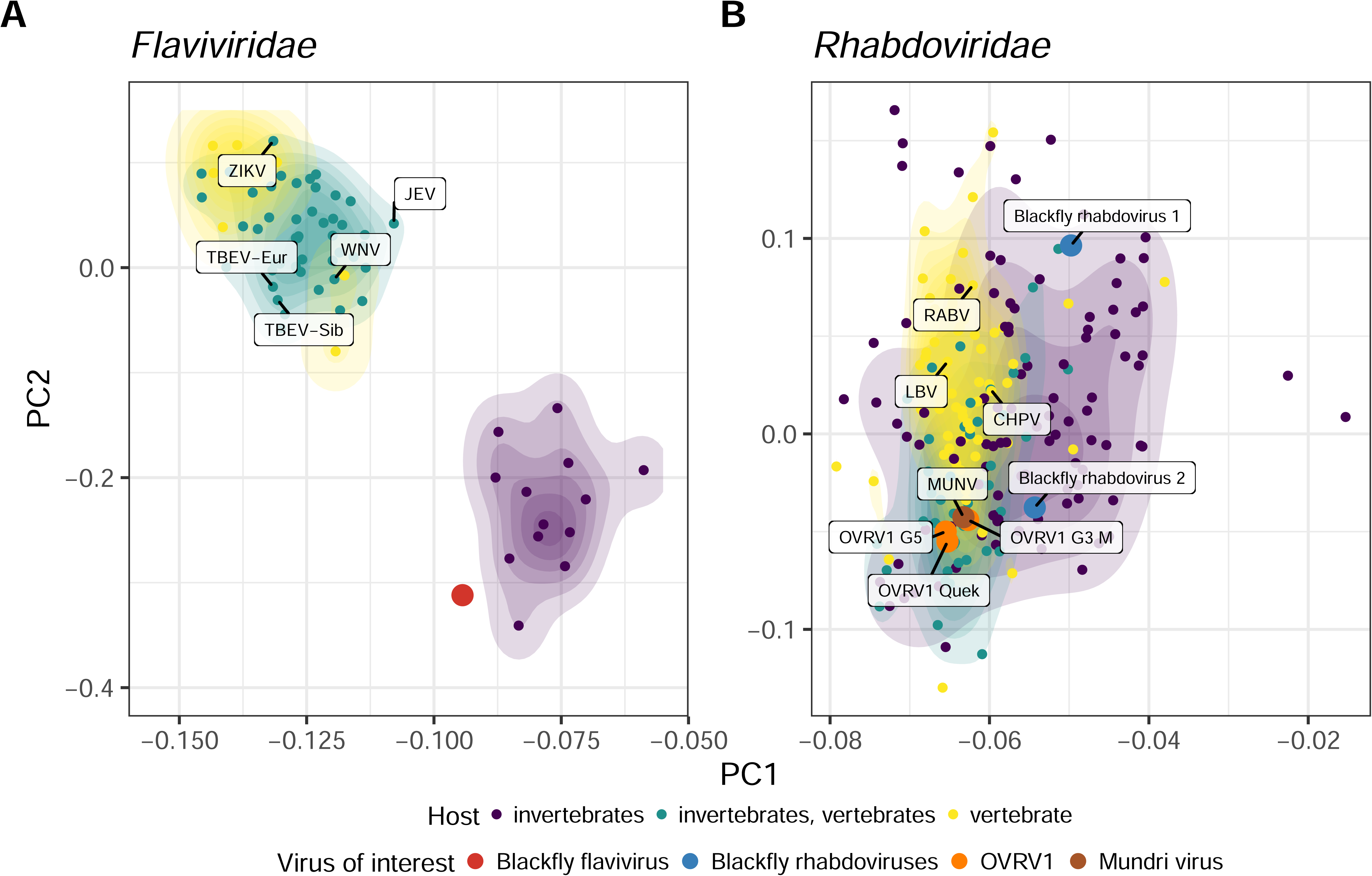
Identified blackfly flavi- and rhabdoviruses are most likely insect-specific. A) PCA plot based on the RDA of selected *Flaviviridae* sequences. The red dot shows the newly identified flavivirus in blackflies. Labels show a selection of neurotropic flaviviruses (TBEV=tick-borne encephalitis virus, JEV=Japanese encephalitis virus, WNV=West Nile virus, ZIKV=Zika virus). B) PCA plot based on the RDA of selected *Rhabdoviridae* sequences. The blue dots show the newly identified blackfly rhabdoviruses, the orange dots show different variants of OVRV1, the brown dot shows Mundri virus (MUNV). Other labelled dots show a selection of neurotropic rhabdoviruses (CHPV=Chandipura virus, LBV=Lagos bat virus, RABV=rabies virus).

## Discussion

To our knowledge, this study represents the first comprehensive investigation of DNA and RNA viruses in blackflies utilizing a whole-genome sequencing approach. A very high number of virus genomes, with a dominance of RNA viruses, were identified in blackfly samples. Fifteen good quality (>50% complete) genomes of eukaryotic DNA viruses (eg. *Parvoviridae* and *Genomoviridae*), and 641 RNA viruses (based on the RdRP, of which 466 with a complete palmprint) were detected. A significant proportion of the novel viruses belonged to viral families associated with human disease, eg. *Flaviviridae, Hepeviridae, Orthomyxoviridae, Parvoviridae, Phenuiviridae, Picornaviridae, Spinareoviridae, Rhabdoviridae*, and *Togaviridae*. However, no viral genomes closely related to known neurotropic viruses were found in the blackfly virome (Supplementary files). Obviously, this does not mean that none of the newly identified viruses could be arboviruses with a neurotropic potential. Additionally, the real host origin for many of the newly discovered viruses remains unknown and further investigations are necessary to determine whether these viruses have the potential to replicate in vertebrate and/or invertebrate hosts, or even if they have a prokaryotic host as could be the case for the *Partitiviridae* and *Picobirnaviridae* (Neri *et al*. 2022; Sadiq, Holmes and Mahar 2024).

The only *Flaviviridae* virus in our analysis was observed at a relatively high abundance in most blackfly samples (see Figure 2). Previous phylogenetic analyses and experimental studies (Kuno and Chang 2005; Cook *et al*. 2013) have shown that the phylogenetic tree of the *Flaviviridae* separates into two large clusters. The first cluster contains ISVs which appear to be host-restricted and only replicate in invertebrate cells and primarily infect mosquitoes (Hoshino *et al*. 2009; Guzman *et al*. 2018; Miranda *et al*. 2019; Baidaliuk *et al*. 2020). The second cluster comprises human pathogenic arboviruses. These previous studies also indicate that arthropod-borne viruses of vertebrates may have originated from ISVs that evolved from being limited to insects to acquiring the ability to infect vertebrate hosts (Bolling *et al*. 2015). The dinucleotide analysis of the *Flaviviridae* indicated that the flavivirus in our blackfly samples is most likely an insect-specific virus.

As blackflies can act as intermediate hosts of pathogens affecting humans and animal health due to their hematophagic habit, several studies (Mead *et al*. 2004; Smith *et al*. 2009; Mesquita *et al*. 2017; Young *et al*. 2021) have focused on the transmission of vesicular stomatitis virus (VSV), an arbovirus belonging to the family *Rhabdoviridae*, genus *Vesiculovirus*, which is known to be transmitted by blackflies. This virus primarily infects livestock, representing an agriculturally relevant pathogen, but zoonotic events have also been reported, highlighting its potential impact on human health (Lichty *et al*. 2004). VSV infections are generally asymptomatic in humans, however, mild flu-like symptoms have been reported in some individuals and a single case of encephalitis in a 3-year-old boy, which was potentially associated with VSV infection, has been reported (Quiroz *et al*. 1988).

In 2022, Edridge and colleagues discovered a divergent rhabdovirus in the bloodstream of a 15-year-old girl with Nodding syndrome from Mundri West Country in South Sudan (Edridge *et al*. 2022). They named this virus “Mundri virus” (MUNV) and classified it as a novel virus. Subsequent phylogenetic analysis revealed that MUNV belongs to the monophyletic clade of *tibroviruses* within the family *Rhabdoviridae*. The *Tibrovirus* genus is part of a larger phylogenetic group of arthropod-borne rhabdoviruses. Several *tibroviruses* have been isolated from biting midges (*Culicoides* spp.) and cattle, and some instances of human detection have been reported (Cybinski *et al*. 1980; Cybinski and Gard 1986; Gibbs *et al*. 1989; Grard *et al*. 2012; Stremlau *et al*. 2015). Human tibrovirus infections may accidentally occur via midge bites (Kuhn *et al*. 2020). Consistent with the findings of Edridge *et al*., our phylogenetic analysis of *Rhabdoviridae* (see Supplemental Figure 13) showed that MUNV is closely related to the *Tibrovirus* genus, supported by a high bootstrap value. However, the two sequences detected in the blackfly virome were not closely related to the *Tibrovirus* nor *Vesiculovirus* genera. In addition, another novel rhabdovirus, named OVRV1, was discovered by Quek *et al*. in public transcriptomes and viromes of *Onchocerca volvulus* nematodes (Quek *et al*. 2024). Because OVRV1 was shown to elicit an immune response in the vertebrate host of the nematode, it could potentially play a role in the pathogenesis of OAE (Quek *et al*. 2024). We were, however, not able to find this virus back in the virome of the blackflies. Our dinucleotide analysis for the *Rhabdoviridae*, suggested that the viruses we found in the blackflies are invertebrate-specific viruses, in contrast to the previously identified rhabdoviruses MUNV (Edridge *et al*. 2022) and OVRV1 (Quek *et al*. 2024). However, a recent association between MUNV infection and nodding syndrome was investigated in a case-control study in South Sudan. MUNV RNA was only detected in the plasma of one of the 72 persons with Nodding syndrome and in none of the controls. MUNV anti-nucleocapsid IgG antibodies were observed in 28% of Nodding syndrome cases, 22% of household controls, and 16% of community controls without significant differences between cases and both control groups. Therefore, it was concluded that MUNV may not be causally associated with Nodding syndrome (Edridge *et al*. 2022). Similarly, it remains to be shown whether OVRV1 may be associated with OAE.

Our study is the first to extensively study the DNA and RNA virome of blackflies from an onchocerciasis-endemic area in Africa. Our findings reveal a relative high abundance of the family *Genomoviridae* in the majority of blackfly samples, aligning with the results of Kraberger *et al*., which demonstrated that the majority of novel ssDNA viral genomes detected in blackflies in New Zealand belonged to the *Genomoviridae* family (Kraberger *et al*. 2019). However, in contrast with the blackflies in New Zealand (Kraberger *et al*. 2019), our study identified *Parvoviridae*, in the blackfly virome. Although this viral family exhibited a high abundance, it was only observed in a limited number of blackfly samples. Phylogenetic analysis revealed that one of the *Parvoviridae* genomes detected in this study clustered within the *Densovirinae* subfamily (see Supplemental Figure 15).

Most of the detected viruses exhibited low sequence similarities with their closest relatives. Consequently, based on the demarcation criteria established for viral classification, it is reasonable to propose that the majority of sequences identified in our study represent new virus species, genera, and/or families, which warrant further studies. In our project, plant and fungal infecting viruses (mainly members of the *Chrysoviridae* and *Partitiviridae* families) were the most abundant, possibly from the nectar diet of the blackflies. Some members of viral families of clinical interest were detected, albeit at lower abundances and no viral genomes closely related to neurotropic viruses were identified.

Our study has some limitations. First of all, it reports the composition of the blackfly virome, including both DNA and RNA viruses but without metadata that would have allowed further associations to be made with *O. volvulus* and potential environmental factors. Further research should compare the virome of blackflies with and without *O. volvulus* infection, and the sex and species of the blackflies should be taken into consideration. It is important to note that only a few blackfly species consume a blood meal (Kraberger *et al*. 2019). Additionally, as less than 1% of blackflies in Cameroon are currently infected with *O. volvulus* (Shintouo *et al*. 2020), it is recommended to screen blackflies for *O. volvulus* infection before sequencing data analyses, as previously suggested (Ekangouo *et al*. 2022).

## Conclusion

Overall, we provide a glimpse into the viral diversity associated with blackfly vectors in Cameroon. Although no genomes associated with neurotropic viruses were found, a plethora of new virus species, genera, and families were identified. The presence of dominant and under-represented viruses warrants future research to determine the role of these viruses in their hosts and ecosystems. Exploring the diversity of viruses in blackflies should be included in the active surveillance of zoonotic diseases. Our findings constitute a starting point for investigating the viruses associated with the hematophagous blackfly and potentially in their nematode host *O. volvulus*.

## Data availability

Sequencing data are available in the SRA database under BioProject accession number PRJNA1088476. Identified viral genomes can be found in following Zenodo repository with a corresponding taxonomy table: https://doi.org/10.5281/zenodo.13737161. The code used for this study is available in a GitHub repository: https://github.com/Matthijnssenslab/2024_XXX_Blackfly_Virome.

## Acknowledgements

We thank Jill Swinnen (KU Leuven) and Jara Wagemans (KU Leuven) for their assistance during the processing of the blackfly pools for Illumina sequencing.

## References

Babaian A, Edgar R. Ribovirus classification by a polymerase barcode sequence. PeerJ 2022;10:e14055.

Baidaliuk A, Lequime S, Moltini-Conclois I et al. Novel genome sequences of cell-fusing agent virus allow comparison of virus phylogeny with the genetic structure of Aedes aegypti populations. Virus Evol 2020;6:1.

Batson J, Dudas G, Haas-Stapleton E et al. Single mosquito metatranscriptomics identifies vectors, emerging pathogens and reservoirs in one assay. Elife 2021;10, DOI: 10.7554/ELIFE.68353.

Bolger AM, Lohse M, Usadel B. Trimmomatic: a flexible trimmer for Illumina sequence data. Bioinformatics 2014;30:2114–20.

Bolling BG, Weaver SC, Tesh RB et al. Insect-specific virus discovery: Significance for the arbovirus community. In Viruses (Vol 2015;7:9.

Boullé C, Njamnshi AK, Dema F et al. Impact of 19 years of mass drug administration with ivermectin on epilepsy burden in a hyperendemic onchocerciasis area in Cameroon. Parasit Vectors 2019;12, DOI: 10.1186/S13071-019-3345-7.

Brattig NW, Cheke RA, Garms R. Onchocerciasis (river blindness) – more than a century of research and control. Acta Trop 2021;218:7.

Buchfink B, Xie C, Huson DH. Fast and sensitive protein alignment using DIAMOND. Nature Methods 2014 12:1 2014;12:59–60.

Camargo AP, Roux S, Schulz F et al. Identification of mobile genetic elements with geNomad. Nature Biotechnology 2023 2023:1–10.

Chesnais CB, Bizet C, Campillo JT et al. A Second Population-Based Cohort Study in Cameroon Confirms the Temporal Relationship Between Onchocerciasis and Epilepsy. Open Forum Infect Dis 2020;7:6.

Chesnais CB, Nana-Djeunga HC, Njamnshi AK et al. The temporal relationship between onchocerciasis and epilepsy: a population-based cohort study. Lancet Infect Dis 2018;18:1278–86.

Colebunders R, Abd-Elfarag G, Carter JY et al. Clinical characteristics of onchocerciasis-associated epilepsy in villages in Maridi County, Republic of South Sudan. Seizure 2018a;62:108–15.

Colebunders R, Hendy A, Nanyunja M et al. Nodding syndrome-a new hypothesis and new direction for research. International Journal of Infectious Diseases 2014;27:74–7.

Colebunders R, Njamnshi AK, Menon S et al. Onchocerca volvulus and epilepsy: A comprehensive review using the Bradford Hill criteria for causation. PLoS Negl Trop Dis 2021;15:1.

Colebunders R, Siewe Fodjo JN, Hopkins A et al. From river blindness to river epilepsy: Implications for onchocerciasis elimination programmes. PLoS Negl Trop Dis 2019;13:7.

Colebunders R, Y. Carter J, Olore PC et al. High prevalence of onchocerciasis-associated epilepsy in villages in Maridi County, Republic of South Sudan: A community-based survey. Seizure 2018b;63:93.

Conceição-Neto N, Zeller M, Lefrère H et al. Modular approach to customise sample preparation procedures for viral metagenomics: A reproducible protocol for virome analysis. Sci Rep 2015;5, DOI: 10.1038/SREP16532.

De Coninck L. ViPER. 2021, DOI: 10.5281/zenodo.5502204.

Cook S, Chung BYW, Bass D et al. Novel virus discovery and genome reconstruction from field rna samples reveals highly divergent viruses in dipteran hosts. PLoS One 2013;8:11.

Van Cutsem G, Siewe Fodjo JN, Dekker MCJ et al. Case definitions for onchocerciasis-associated epilepsy and nodding syndrome: A focused review. Seizure 2023;107:132–5.

Cybinski DH, Gard GP. Isolation of a new rhabdovirus in Australia related to Tibrogargan virus. Aust J Biol Sci 1986;39:225–32.

Cybinski DH, St. George TD, Standfast HA et al. Isolation of tibrogargan virus, a new Australian rhabdovirus, from Culicoides brevitarsis. Vet Microbiol 1980;5:301–8.

Davis NM, Proctor DiM, Holmes SP et al. Simple statistical identification and removal of contaminant sequences in marker-gene and metagenomics data. Microbiome 2018;6:1–14.

Dudas G, Batson J. Accumulated metagenomic studies reveal recent migration, whole genome evolution, and undiscovered diversity of orthomyxoviruses. J Virol 2023;97, DOI: 10.1128/JVI.01056-23/SUPPL_FILE/JVI.01056-23-S0001.PDF.

Duke BO, Vincelette J, Moore PJ. Microfilariae in the cerebrospinal fluid, and neurological complications, during treatment of onchocerciasis with diethylcarbamazine. Tropenmed Parasitol 1976;27:123–32.

Eddy SR. Accelerated Profile HMM Searches. PLoS Comput Biol 2011;7, DOI: 10.1371/JOURNAL.PCBI.1002195.

Edgar RC. Muscle5: High-accuracy alignment ensembles enable unbiased assessments of sequence homology and phylogeny. Nature Communications 2022 13:1 2022;13:1–9.

Edridge AWD, Abd-Elfarag G, Deijs M et al. Divergent Rhabdovirus Discovered in a Patient with New-Onset Nodding Syndrome. Viruses 2022;14:210.

Ekangouo AE, Djeunga HCN, Sempere G et al. Bacteriome Diversity of Blackflies’ Gut and Association with Onchocerca volvulus, the Causative Agent of Onchocerciasis in Mbam Valley (Center Region, Cameroon). Pathogens 2022;11, DOI: 10.3390/PATHOGENS11010044.

Föger K, Gora-Stahlberg G, Sejvar J et al. Nakalanga Syndrome: Clinical Characteristics, Potential Causes, and Its Relationship with Recently Described Nodding Syndrome. PLoS Negl Trop Dis 2017;11, DOI: 10.1371/JOURNAL.PNTD.0005201.

Fros JJ, Visser I, Tang B et al. The dinucleotide composition of the Zika virus genome is shaped by conflicting evolutionary pressures in mammalian hosts and mosquito vectors. PLoS Biol 2021;19, DOI: 10.1371/JOURNAL.PBIO.3001201.

Gibbs EPJ, Calisher CH, Tesh RB et al. Bivens arm virus: a new rhabdovirus isolated from Culicoides insignis in Florida and related to Tibrogargan virus of Australia. Vet Microbiol 1989;19:141–50.

Grard G, Fair JN, Lee D et al. A novel rhabdovirus associated with acute hemorrhagic fever in central Africa. PLoS Pathog 2012;8, DOI: 10.1371/JOURNAL.PPAT.1002924.

Guzman H, Contreras-Gutierrez MA, Travassos da Rosa APA et al. Characterization of Three New Insect-Specific Flaviviruses: Their Relationship to the Mosquito-Borne Flavivirus Pathogens. Am J Trop Med Hyg 2018;98:410.

Hadermann A, Amaral LJ, Van Cutsem G et al. Onchocerciasis-associated epilepsy: an update and future perspectives. Trends Parasitol 2023;39:126–38.

Hendy A, Krit M, Pfarr K et al. Onchocerca volvulus transmission in the Mbam valley of Cameroon following 16 years of annual community-directed treatment with ivermectin, and the description of a new cytotype of Simulium squamosum. Parasit Vectors 2021;14:563.

Hoshino K, Isawa H, Tsuda Y et al. Isolation and characterization of a new insect flavivirus from Aedes albopictus and Aedes flavopictus mosquitoes in Japan. Virology 2009;391:1.

Hotterbeekx A, Lammens M, Idro R et al. Neuroinflammation and not tauopathy is a predominant pathological signature of nodding syndrome. J Neuropathol Exp Neurol 2019;78:1049–58.

Hotterbeekx A, Raimon S, Abd-Elfarag G et al. Onchocerca volvulus is not detected in the cerebrospinal fluid of persons with onchocerciasis-associated epilepsy. International Journal of Infectious Diseases 2020;91:119–23.

Howerth EW, Mead DG, Stallknecht DE. Immunolocalization of vesicular stomatitis virus in black flies (Simulium vittatum). Ann N Y Acad Sci 2002;969:340.

Hyatt D, Chen GL, LoCascio PF et al. Prodigal: Prokaryotic gene recognition and translation initiation site identification. BMC Bioinformatics 2010;11:1–11.

Kraberger S, Schmidlin K, Fontenele RS et al. Unravelling the single-stranded DNA virome of the New Zealand blackfly. Viruses 2019;11, DOI: 10.3390/V11060532.

Kuhn JH, Pan H, Chiu CY et al. Human Tibroviruses: Commensals or Lethal Pathogens? Viruses 2020;12:3.

Kuno G, Chang GJJ. Biological transmission of arboviruses: Reexamination of and new insights into components, mechanisms, and unique traits as well as their evolutionary trends. In Clinical Microbiology Reviews (Vol 2005;18:4.

Levick B, Laudisoit A, Tepage F et al. High prevalence of epilepsy in onchocerciasis endemic regions in the Democratic Republic of the Congo. PLoS Negl Trop Dis 2017;11:e0005732.

Lichty BD, Power AT, Stojdl DF et al. Vesicular stomatitis virus: Re-inventing the bullet. Trends Mol Med 2004;10:5.

Lytras S, Hughes J. Synonymous Dinucleotide Usage: A Codon-Aware Metric for Quantifying Dinucleotide Representation in Viruses. Viruses 2020, Vol 12, Page 462 2020;12:462.

Mead DG, Howerth EW, Murphy MD et al. Black fly involvement in the epidemic transmission of Vesicular stomatitis New Jersey virus (Rhabdoviridae: Vesiculovirus). Vector-Borne and Zoonotic Diseases 2004;4:351–9.

Mesquita LP, Diaz MH, Howerth EW et al. Pathogenesis of Vesicular Stomatitis New Jersey Virus Infection in Deer Mice (Peromyscus maniculatus) Transmitted by Black Flies (Simulium vittatum). Vet Pathol 2017;54:1.

Miranda J, Mattar S, Gonzalez M et al. First report of Culex flavivirus infection from Culex coronator (Diptera: Culicidae), Colombia. Virol J 2019;16, DOI: 10.1186/S12985-018-1108-2.

Mukendi D, Tepage F, Akonda I et al. High prevalence of epilepsy in an onchocerciasis endemic health zone in the Democratic Republic of the Congo, despite 14 years of community-directed treatment with ivermectin: A mixed-method assessment. Int J Infect Dis 2019;79:187–94.

Nayfach S, Camargo AP, Schulz F et al. CheckV assesses the quality and completeness of metagenome-assembled viral genomes. Nature Biotechnology 2020 39:5 2020;39:578–85.

Neri U, Wolf YI, Roux S et al. Expansion of the global RNA virome reveals diverse clades of bacteriophages. Cell 2022;185:4023–4037.e18.

Nurk S, Meleshko D, Korobeynikov A et al. MetaSPAdes: A new versatile metagenomic assembler. Genome Res 2017;27:824–34.

Piégu B, Guizard S, Spears T et al. Complete genome sequence of invertebrate iridescent virus 22 isolated from a blackfly larva. Journal of General Virology 2013;94:9.

Piégu B, Guizard S, Spears T et al. Complete genome sequence of invertebrate iridovirus IIV-25 isolated from a blackfly larva. Arch Virol 2014a;159:5.

Piégu B, Guizard S, Yeping T et al. Complete genome sequence of invertebrate iridovirus IIV22A, a variant of IIV22, isolated originally from a blackfly larva. Stand Genomic Sci 2014b;9:3.

Pion SDS, Boussinesq M. Significant Association between Epilepsy and Presence of Onchocercal Nodules: Case-Control Study in Cameroon. Am J Trop Med Hyg 2012;86:557.

Price MN, Dehal PS, Arkin AP. FastTree 2 – Approximately Maximum-Likelihood Trees for Large Alignments. PLoS One 2010;5, DOI: 10.1371/JOURNAL.PONE.0009490.

Quek S, Hadermann A, Wu Y et al. Diverse RNA viruses of parasitic nematodes can elicit antibody responses in vertebrate hosts. Nature Microbiology 2024 2024:1–18.

Quiroz E, Moreno N, Peralta PH et al. A human case of encephalitis associated with vesicular stomatitis virus (Indiana serotype) infection. Am J Trop Med Hyg 1988;39:3.

Raimon S, Dusabimana A, Abd-Elfarag G et al. High Prevalence of Epilepsy in an Onchocerciasis-Endemic Area in Mvolo County, South Sudan: A Door-To-Door Survey. Pathogens 2021;10, DOI: 10.3390/PATHOGENS10050599.

Sadiq S, Holmes EC, Mahar JE. Genomic and phylogenetic features of the Picobirnaviridae suggest microbial rather than animal hosts. bioRxiv 2024:2024.02.04.578841.

Sakaguchi S, Urayama SI, Takaki Y et al. NeoRdRp: A Comprehensive Dataset for Identifying RNA-dependent RNA Polymerases of Various RNA Viruses from Metatranscriptomic Data. Microbes Environ 2022;37, DOI: 10.1264/JSME2.ME22001.

Shintouo CM, Nguve JE, Asa FB et al. Entomological assessment of onchocerca species transmission by black flies in selected communities in the west region of cameroon. Pathogens 2020;9:9.

Siewe Fodjo JN, Mandro M, Mukendi D et al. Onchocerciasis-associated epilepsy in the Democratic Republic of Congo: Clinical description and relationship with microfilarial density. PLoS Negl Trop Dis 2019;13, DOI: 10.1371/JOURNAL.PNTD.0007300.

Siewe Fodjo JN, Ngarka L, Njamnshi WY et al. Onchocerciasis in the Ntui Health District of Cameroon: epidemiological, entomological and parasitological findings in relation to elimination prospects. Parasit Vectors 2022;15, DOI: 10.1186/S13071-022-05585-0.

Siewe Fodjo JN, Tatah G, Tabah EN et al. Epidemiology of onchocerciasis-associated epilepsy in the Mbam and Sanaga river valleys of Cameroon: Impact of more than 13 years of ivermectin. Infect Dis Poverty 2018;7:1.

Smith PF, Howerth EW, Carter D et al. Mechanical transmission of vesicular stomatitis New Jersey virus by Simulium vittatum (Diptera: Simuliidae) to domestic swine (Sus scrofa). J Med Entomol 2009;46:6.

Stremlau MH, Andersen KG, Folarin OA et al. Discovery of novel rhabdoviruses in the blood of healthy individuals from West Africa. PLoS Negl Trop Dis 2015;9, DOI: 10.1371/JOURNAL.PNTD.0003631.

Winkler AS, Friedrich K, Velicheti S et al. MRI findings in people with epilepsy and nodding syndrome in an area endemic for onchocerciasis: An observational study. Afr Health Sci 2013;13:529–40.

Wood DE, Lu J, Langmead B. Improved metagenomic analysis with Kraken 2. Genome Biol 2019;20:1–13.

Young KI, Valdez F, Vaquera C et al. Surveillance along the Rio Grande during the 2020 Vesicular Stomatitis Outbreak Reveals Spatio-Temporal Dynamics of and Viral RNA Detection in Black Flies. Pathogens 2021;10:10.

