## Supplementary Figures and Tables for "Cameroonian blackflies (Diptera: Simuliidae) harbour a plethora of (RNA) viruses"

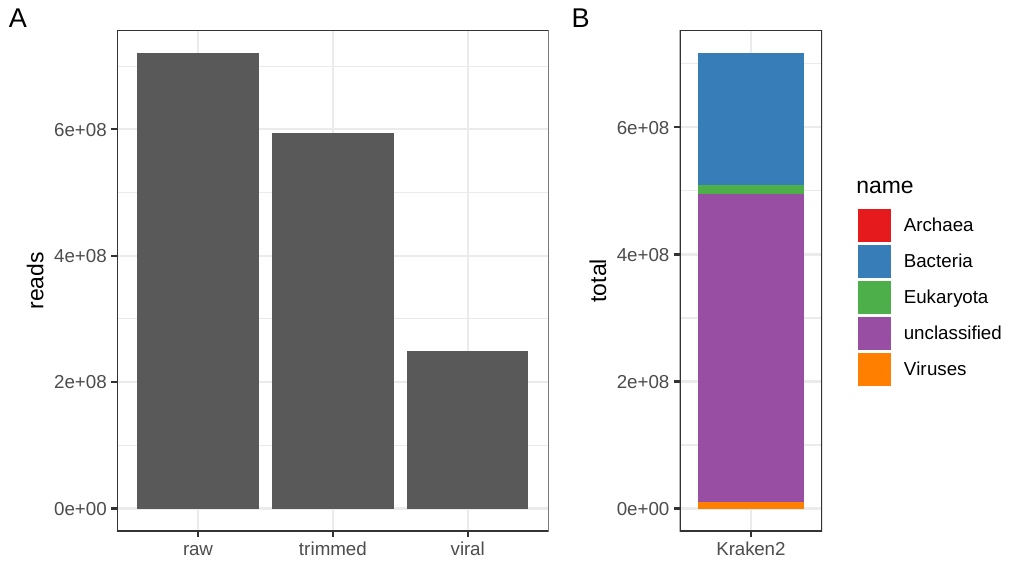

**Supplementary Figure 1**

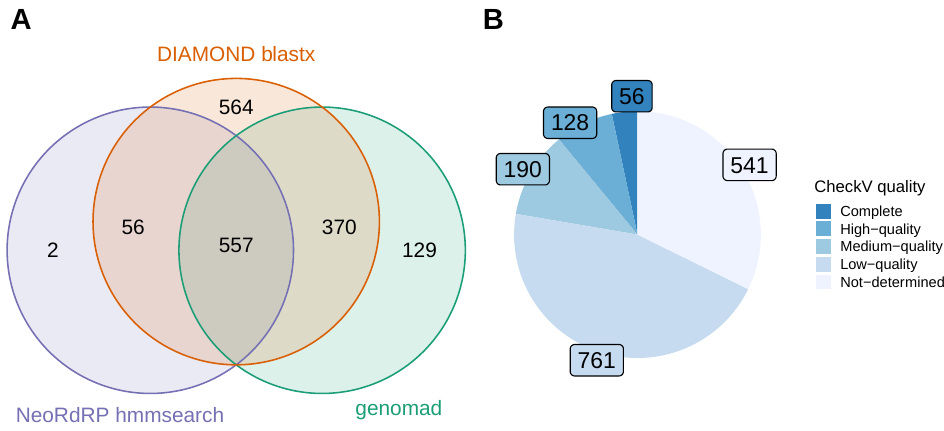

**Supplementary Figure 2**

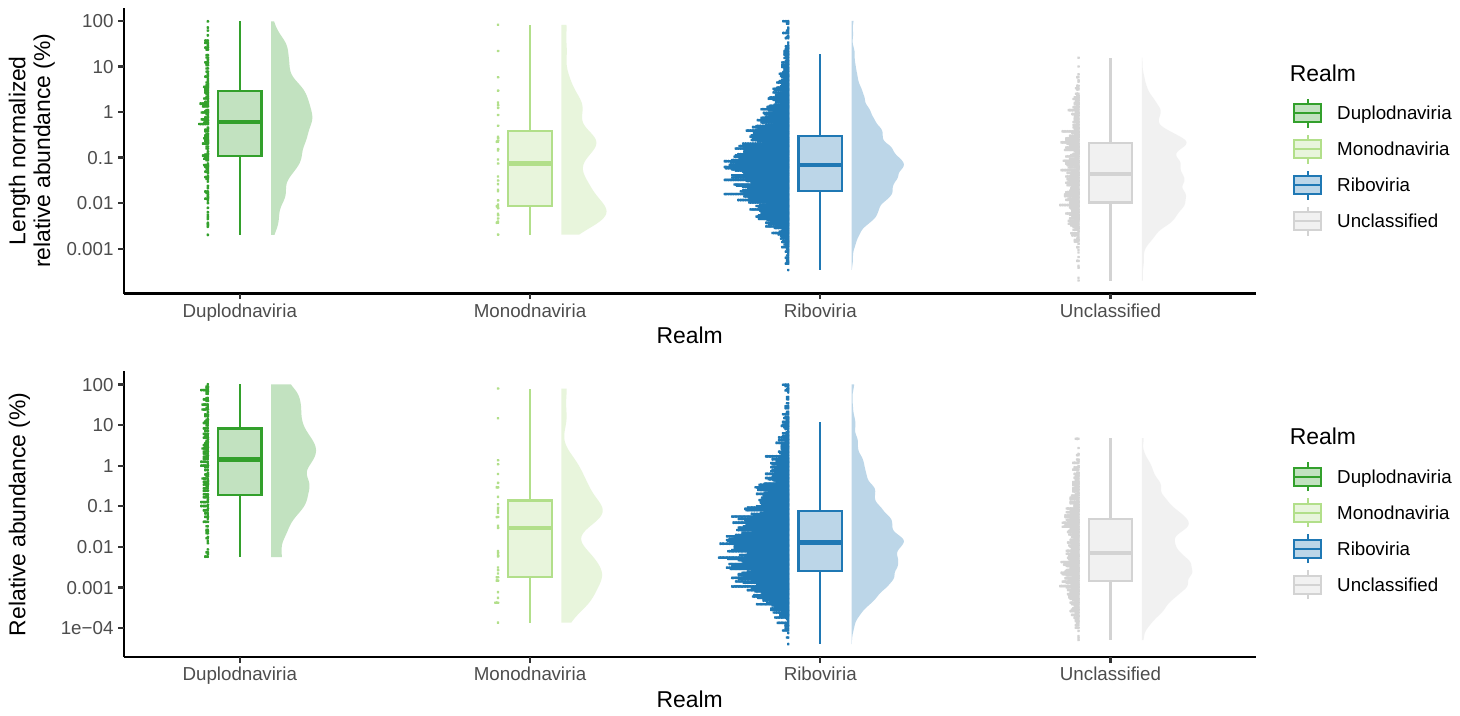

**Supplementary Figure 3**

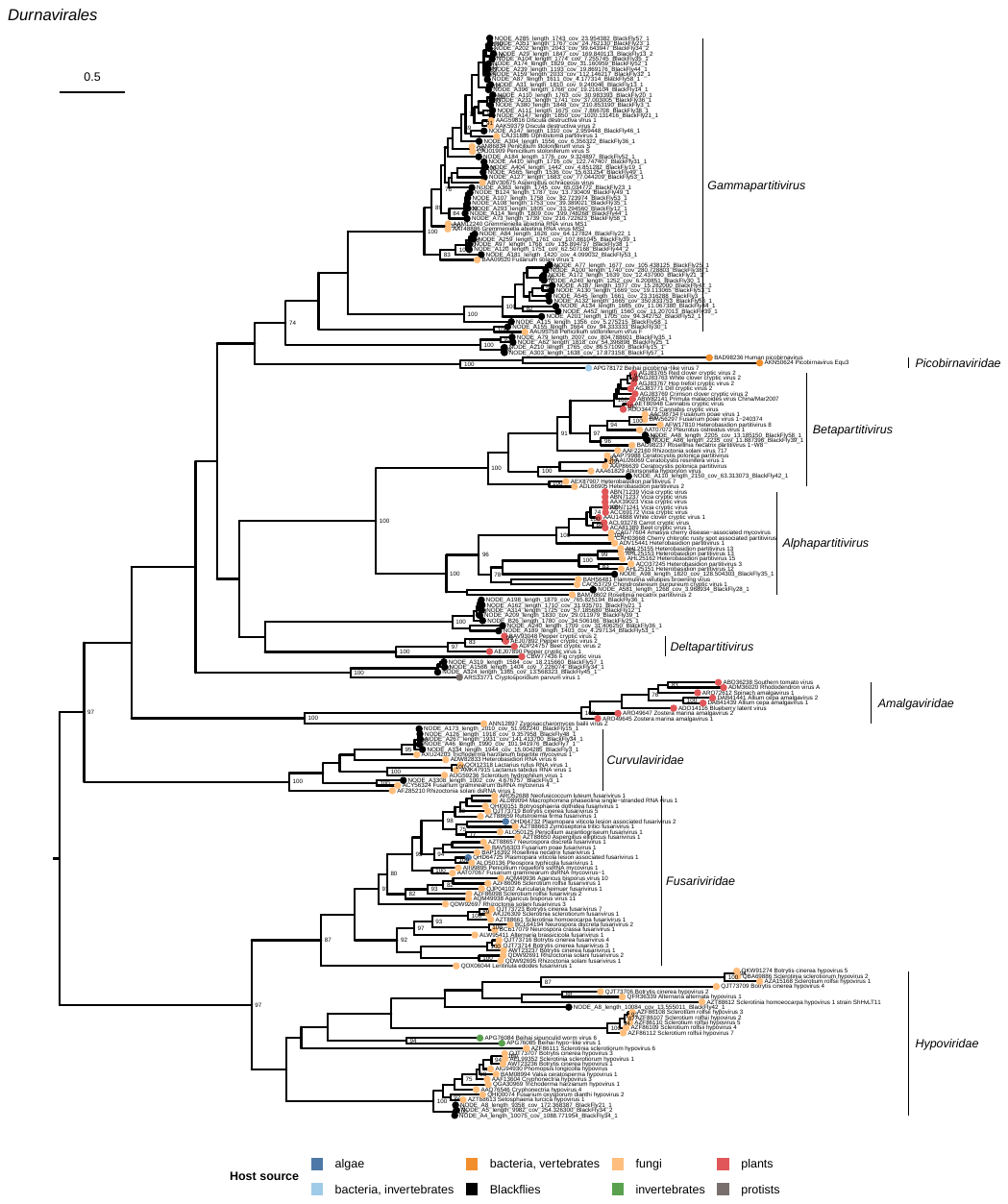

**Supplementary Figure 4**

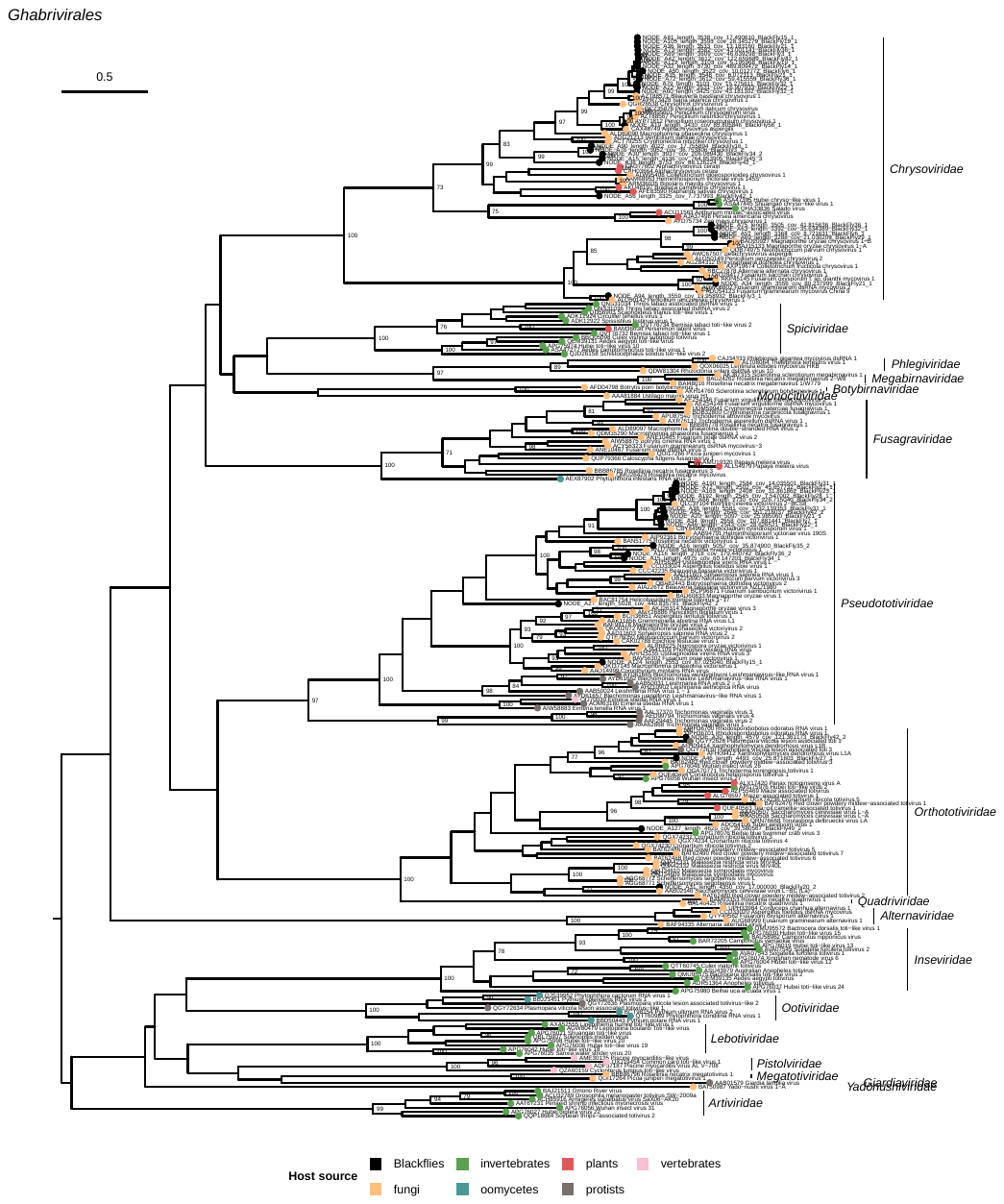

**Supplementary Figure 5**

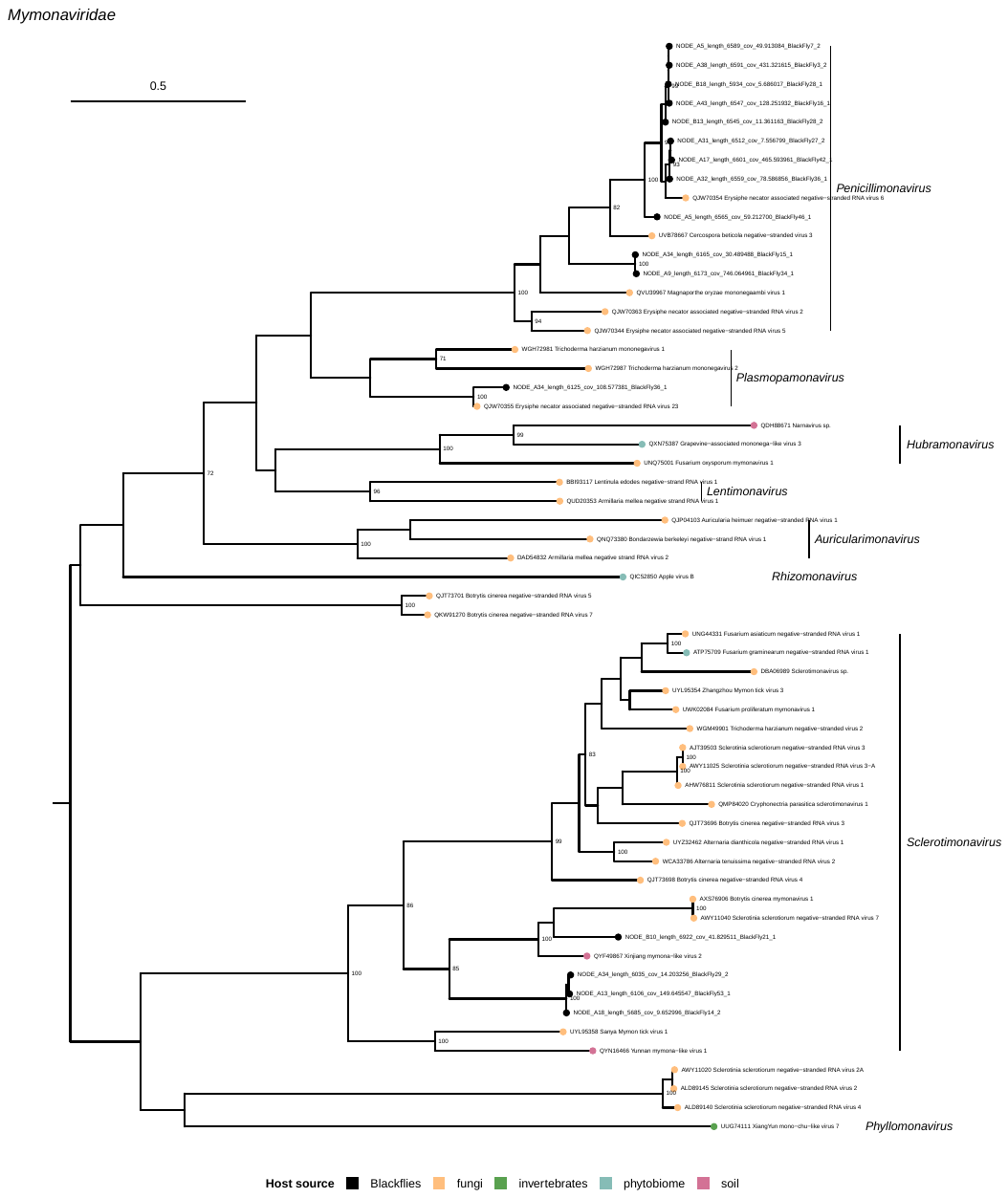

**Supplementary Figure 6**

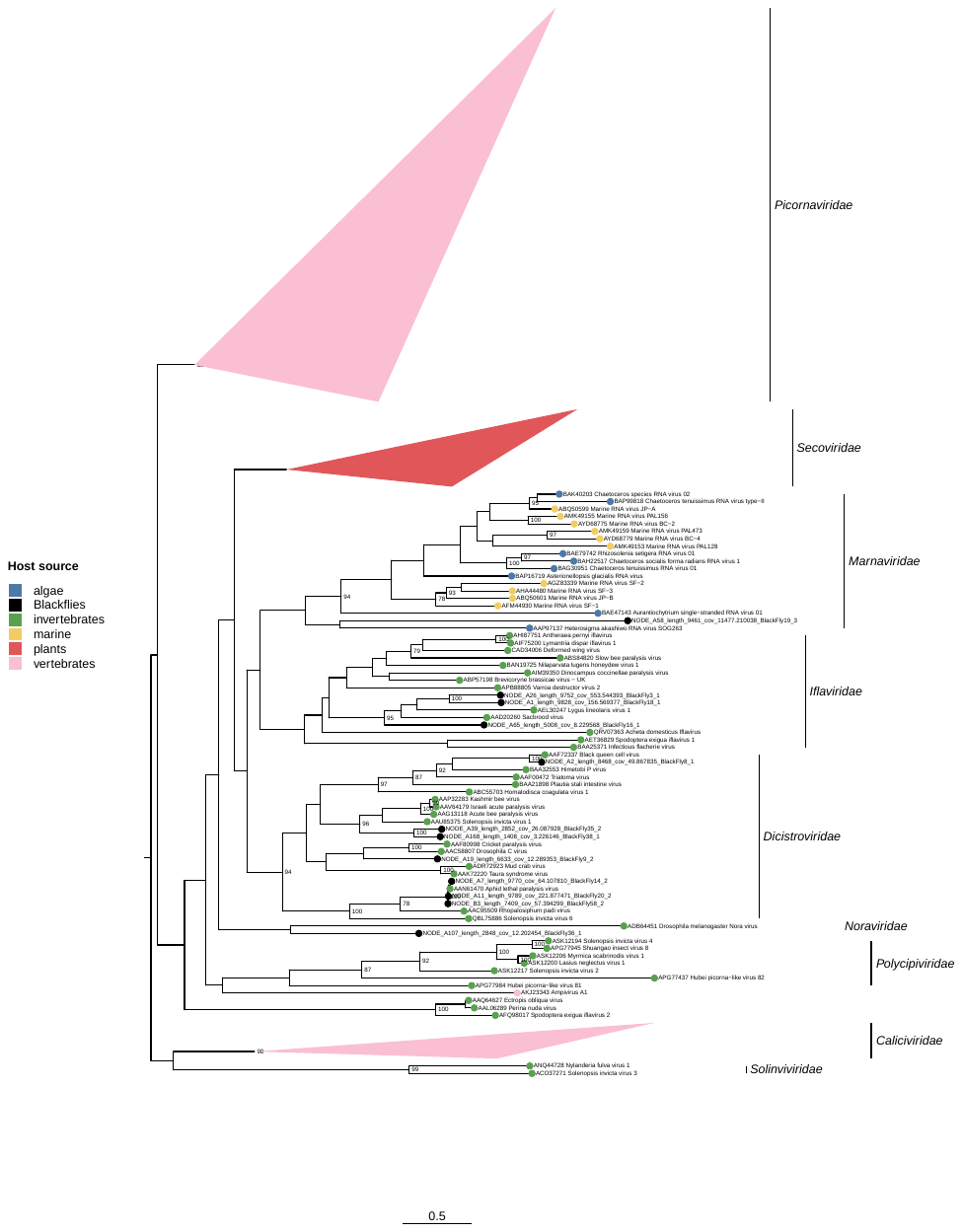

**Supplementary Figure 7**

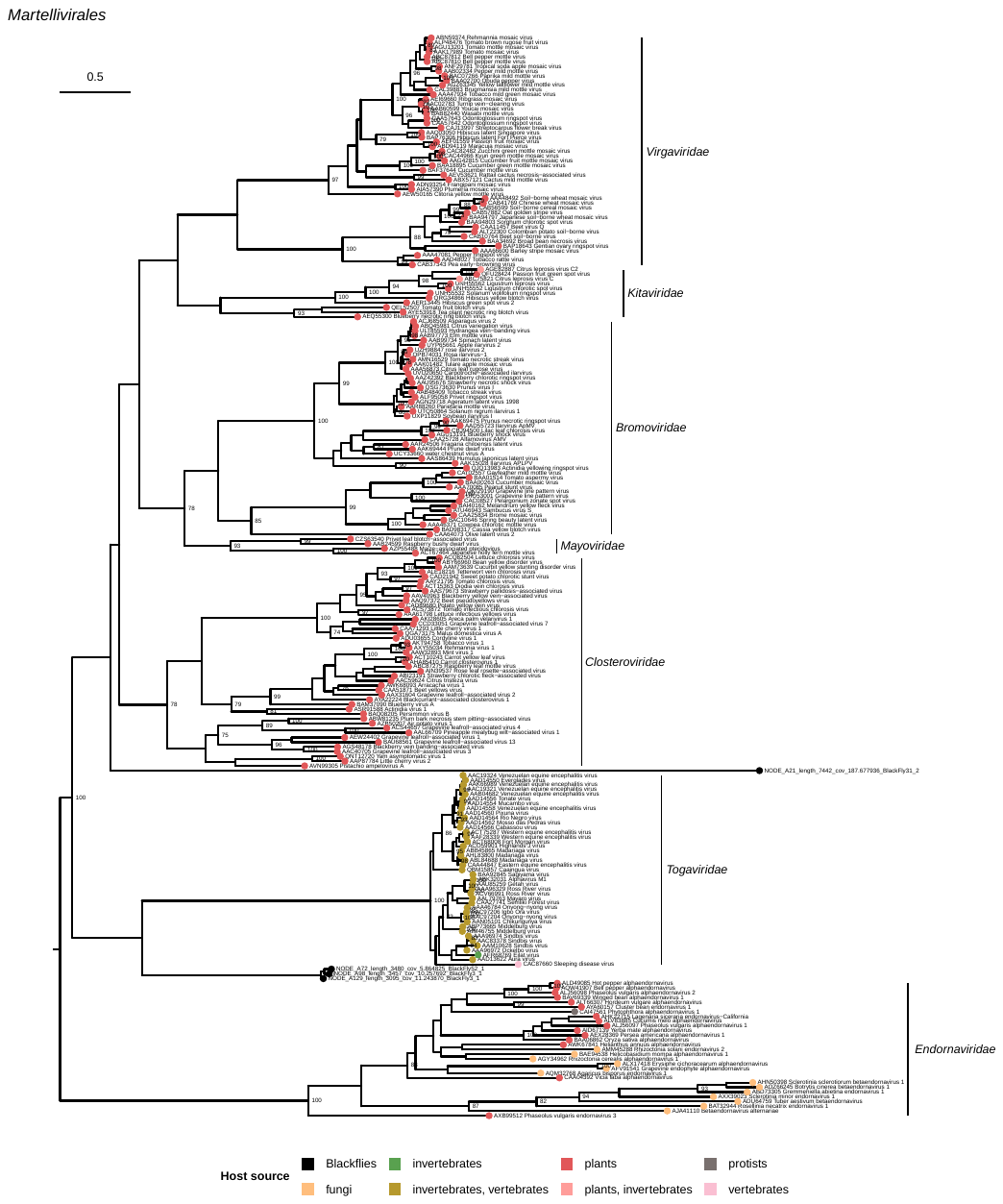

**Supplementary Figure 8**

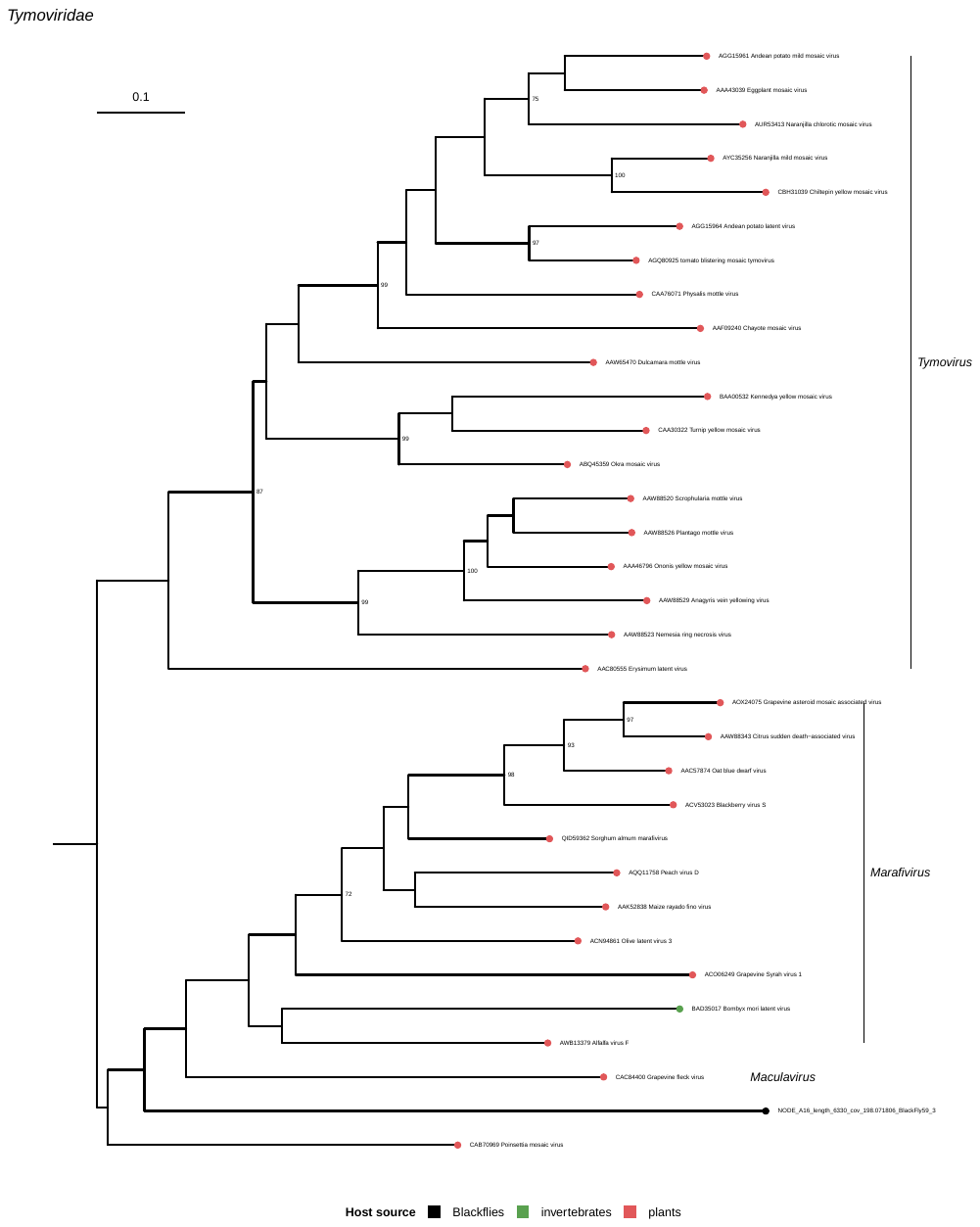

**Supplementary Figure 9**

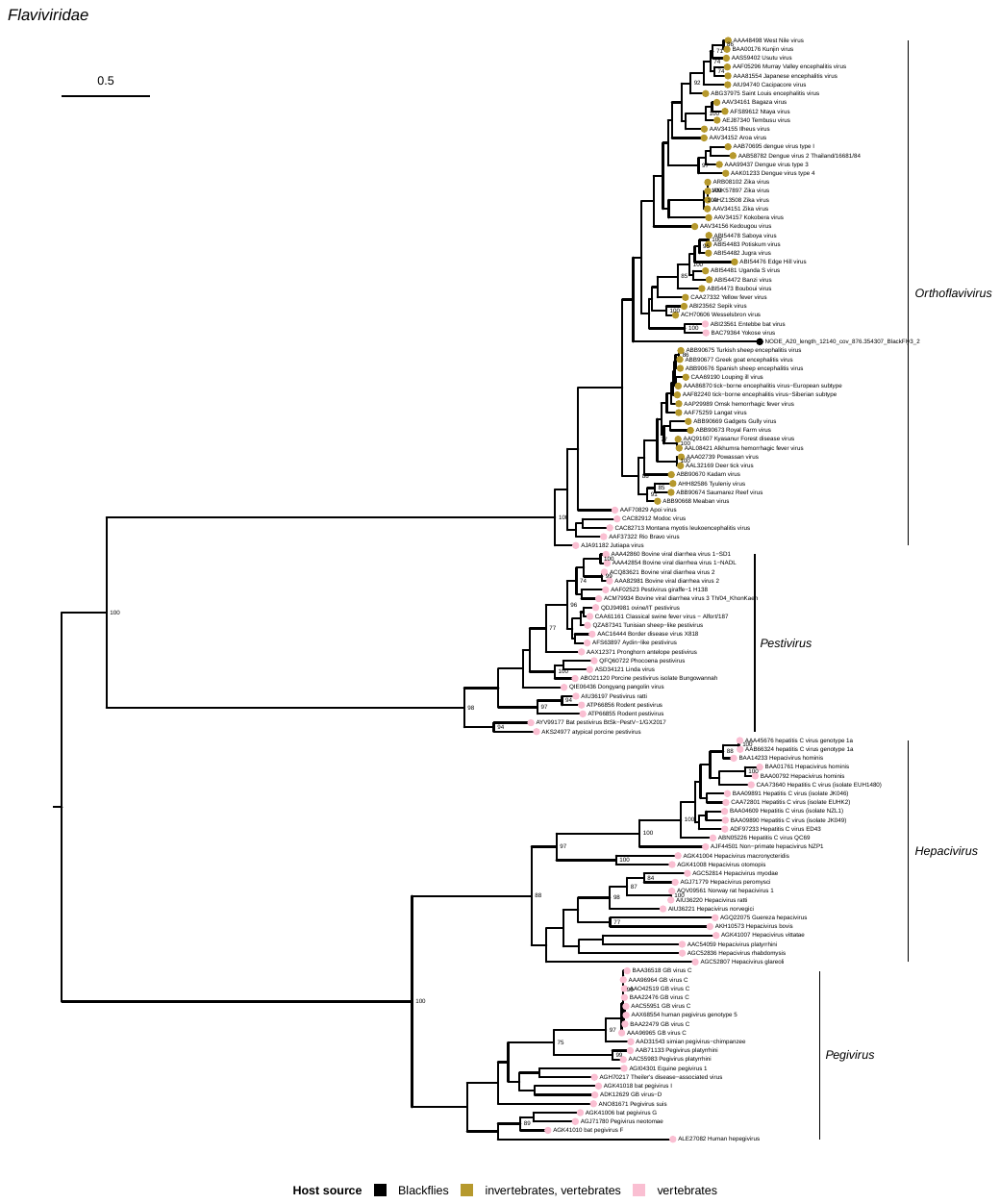

**Supplementary Figure 10**

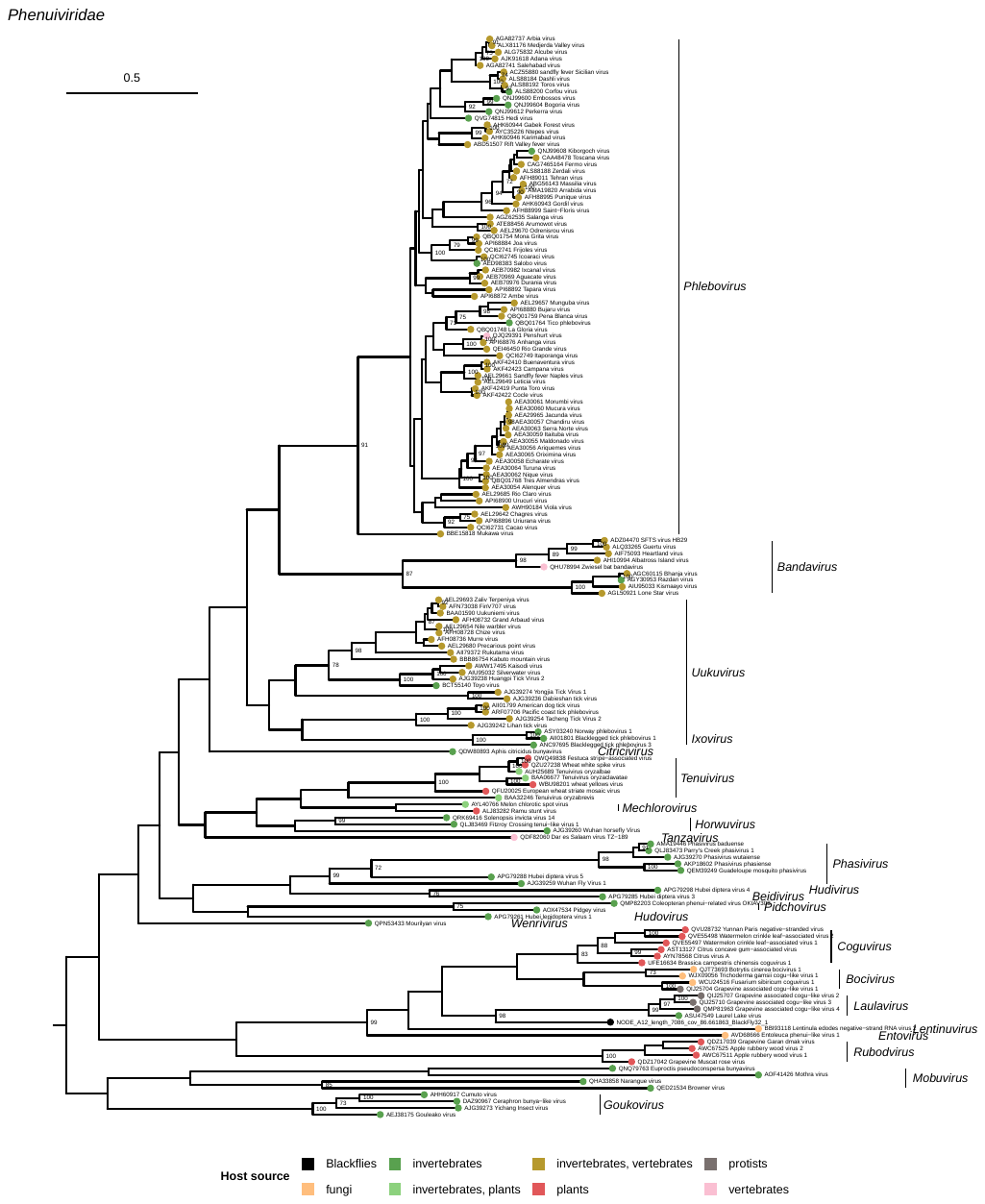

**Supplementary Figure 11**

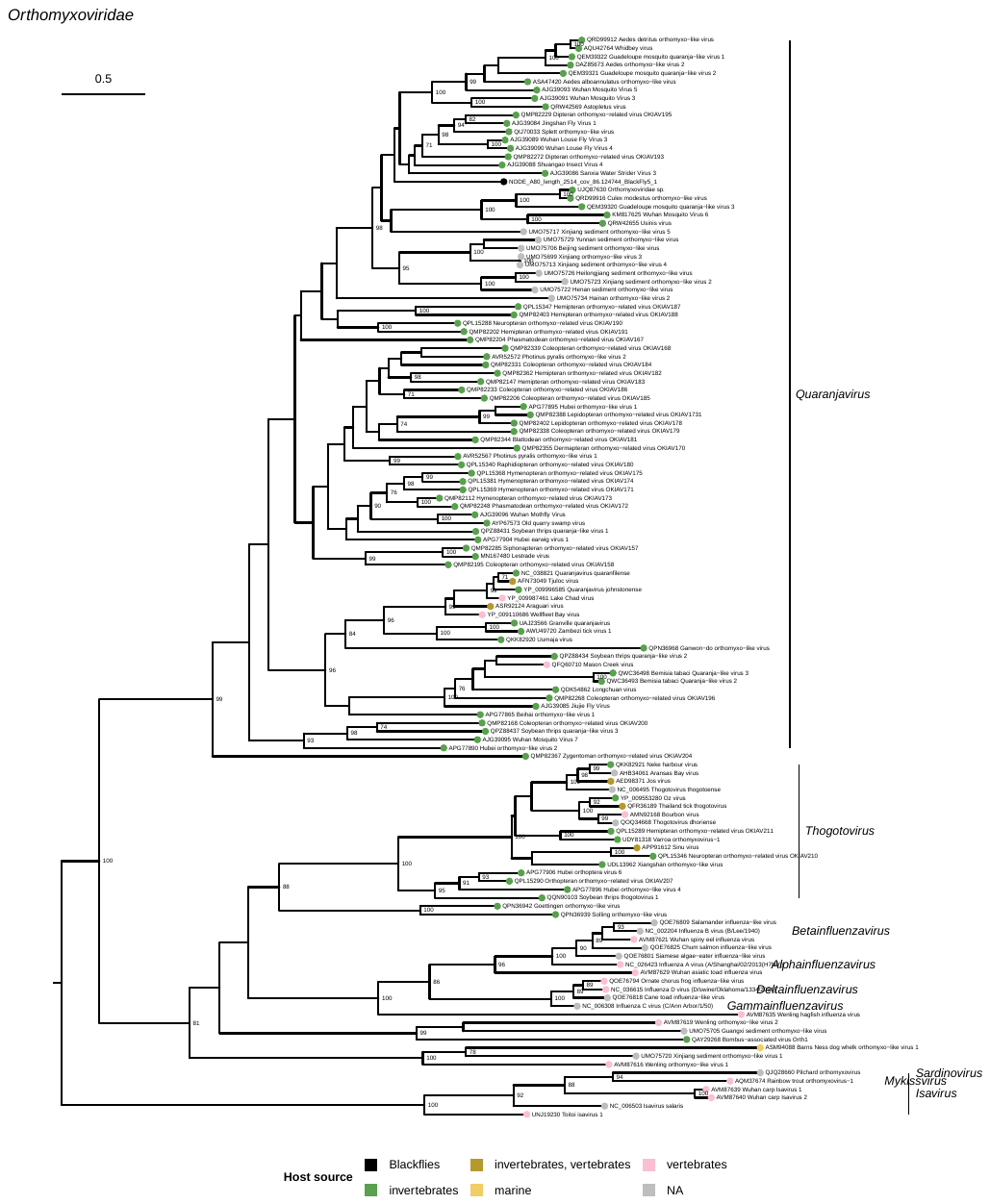

**Supplementary Figure 12**

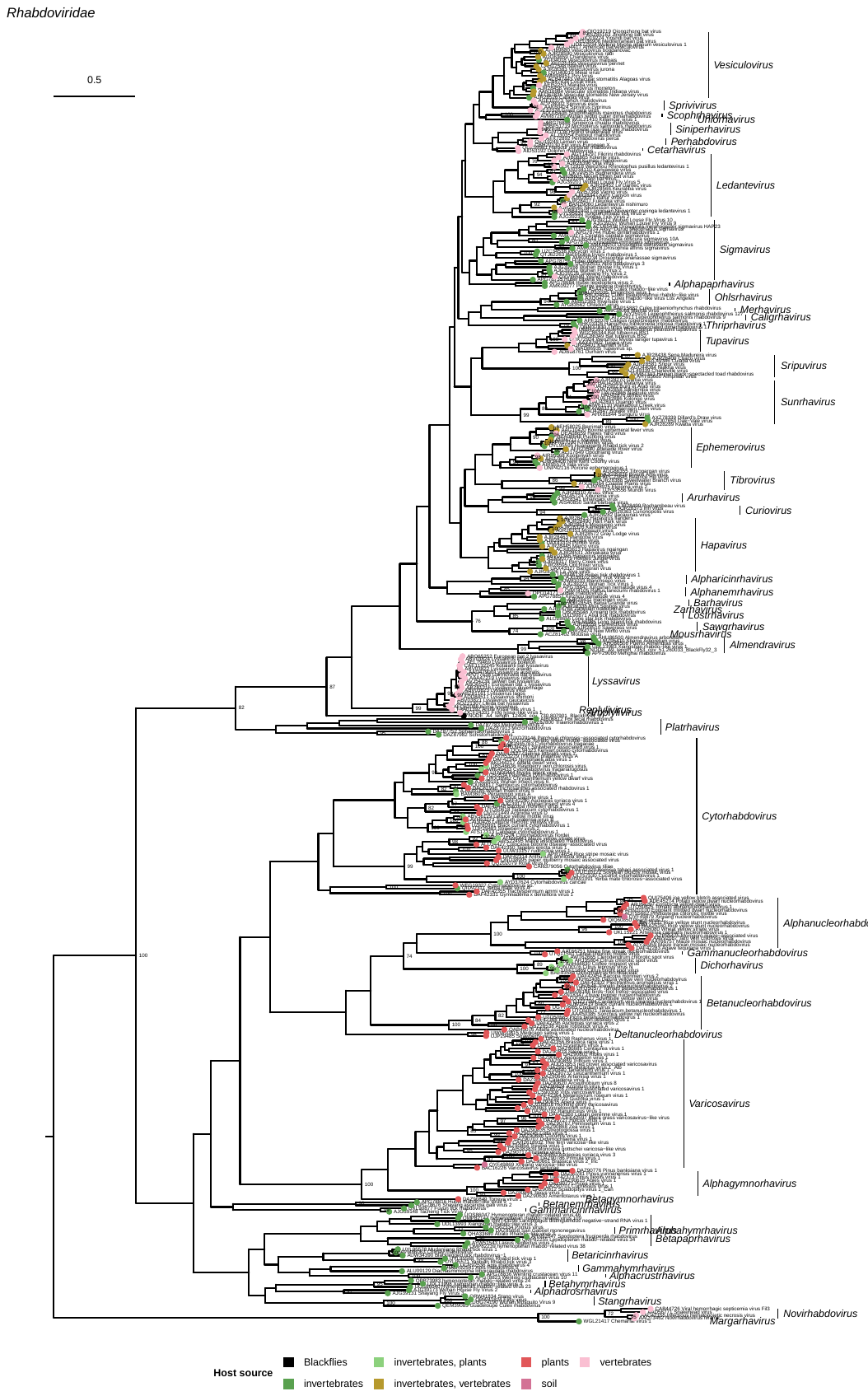
**Supplementary Figure 13**

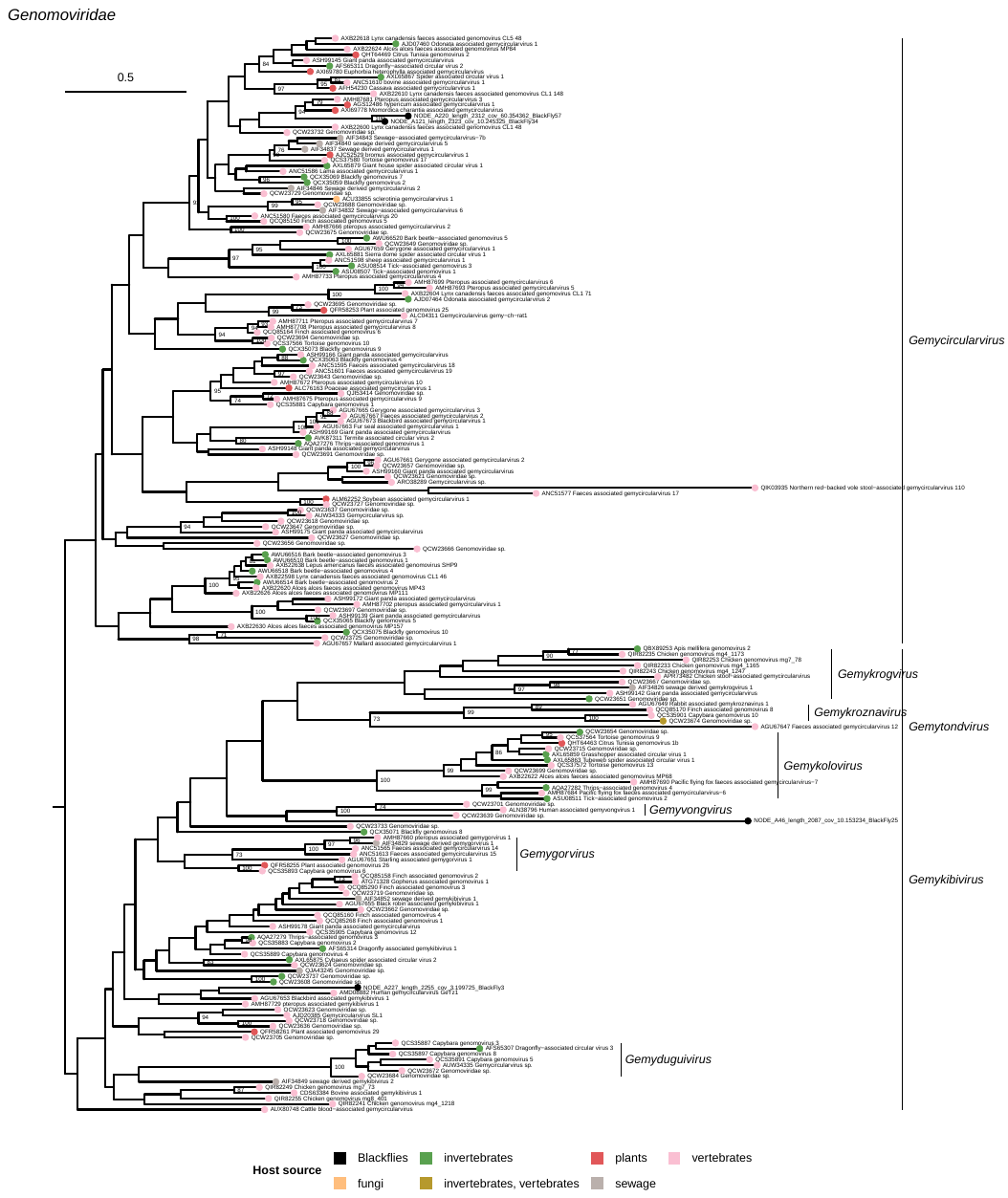

**Supplementary Figure 14**

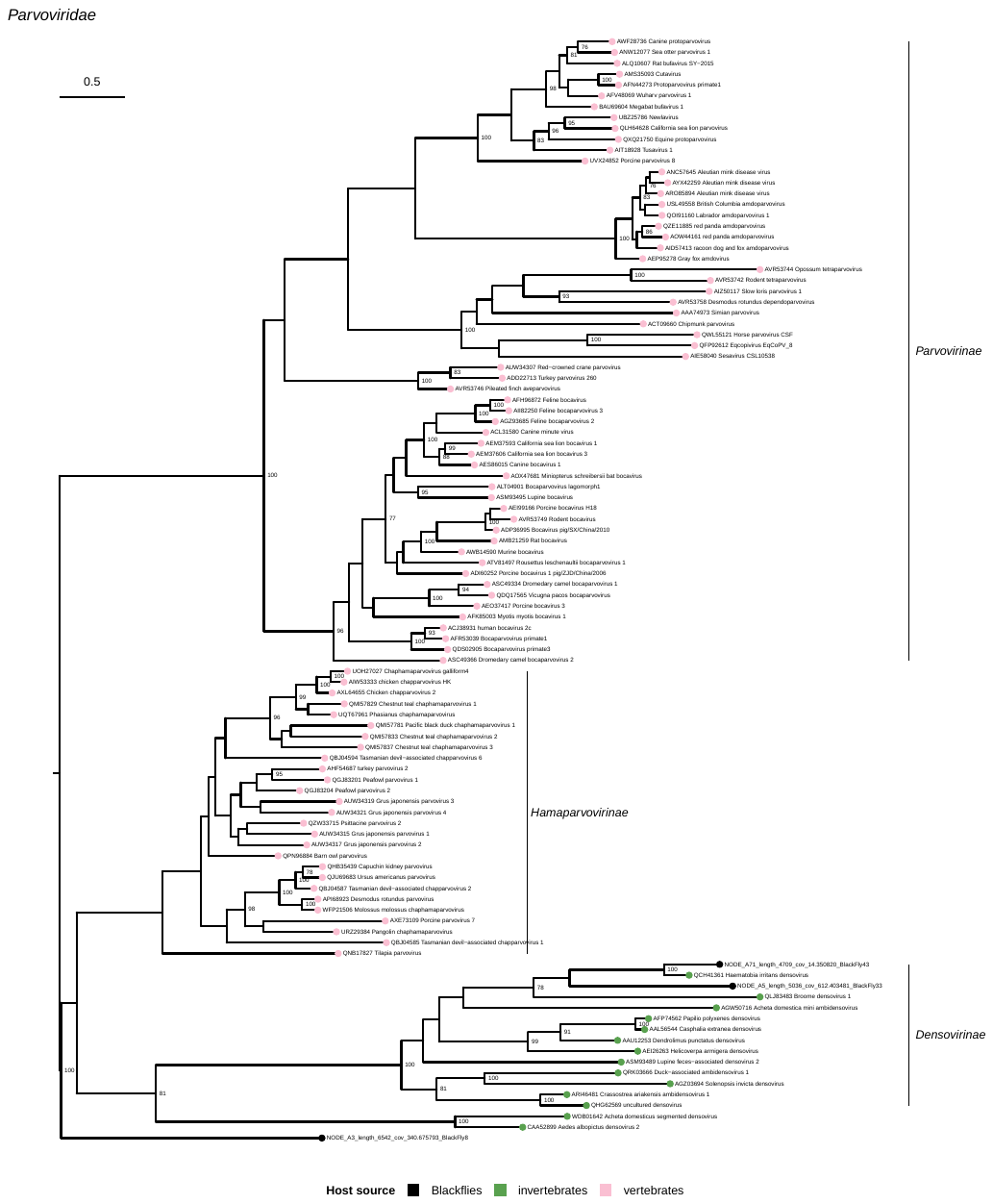

**Supplementary Figure 15**

Supplementary Table 1

| Sample | num_seqs_raw | sum_len_raw | num_seqs_trimmed | sum_len_trimmed |
| --- | --- | --- | --- | --- |
| BlackFly3 | 15967058 | 2411025758 | 14083411 | 1802680773 |
| BlackFly4 | 10618738 | 1603429438 | 9517543 | 1192269936 |
| BlackFly5 | 16150320 | 2438698320 | 14102123 | 1717230119 |
| BlackFly6 | 12647152 | 1909719952 | 11036288 | 1353645999 |
| BlackFly7 | 9975134 | 1506245234 | 8399644 | 1023297094 |
| BlackFly8 | 9969112 | 1505335912 | 7795933 | 931437183 |
| BlackFly9 | 13861100 | 2093026100 | 11036751 | 1253421775 |
| BlackFly10 | 9500948 | 1434643148 | 7377393 | 851224264 |
| BlackFly11 | 11569104 | 1746934704 | 8932398 | 1016081035 |
| BlackFly12 | 17760656 | 2681859056 | 16170603 | 2017817725 |
| BlackFly13 | 11621738 | 1754882438 | 9761262 | 1121529459 |
| BlackFly14 | 13825740 | 2087686740 | 11367218 | 1368627775 |
| BlackFly15 | 12254150 | 1850376650 | 10898369 | 1343794670 |
| BlackFly16 | 23262772 | 3512678572 | 20790370 | 2593968097 |
| BlackFly17 | 8487868 | 1281668068 | 6552560 | 710985522 |
| BlackFly18 | 11315546 | 1708647446 | 8810100 | 975232370 |
| BlackFly19 | 17325276 | 2616116676 | 14611737 | 1783268241 |
| BlackFly20 | 21282790 | 3213701290 | 18469265 | 2287470708 |
| BlackFly21 | 11482716 | 1733890116 | 9342539 | 1068919360 |
| BlackFly22 | 7261050 | 1096418550 | 5272569 | 563885155 |
| BlackFly23 | 18917278 | 2856508978 | 16131017 | 2005350235 |
| BlackFly24 | 9540220 | 1440573220 | 7108174 | 752892879 |
| BlackFly25 | 12024060 | 1815633060 | 9368749 | 1076209165 |
| BlackFly26 | 9754940 | 1472995940 | 7576311 | 859096268 |
| BlackFly27 | 18857860 | 2847536860 | 15629349 | 1884197878 |
| BlackFly28 | 16220006 | 2449220906 | 14396532 | 1753659158 |
| BlackFly29 | 19156540 | 2892637540 | 16901664 | 2088463894 |
| BlackFly30 | 20808714 | 3142115814 | 18463756 | 2341331981 |
| BlackFly31 | 21630972 | 3266276772 | 19174474 | 2436102195 |
| BlackFly32 | 14763992 | 2229362792 | 12656550 | 1548953159 |
| BlackFly33 | 6113870 | 923194370 | 4898036 | 528128022 |
| BlackFly34 | 24002232 | 3624337032 | 20578254 | 2601183937 |
| BlackFly35 | 16871692 | 2547625492 | 14462414 | 1790150100 |
| BlackFly36 | 14992036 | 2263797436 | 12568209 | 1551430241 |
| BlackFly37 | 7467628 | 1127611828 | 4544534 | 440090237 |
| BlackFly38 | 10610832 | 1602235632 | 7890352 | 900521109 |
| BlackFly39 | 13739574 | 2074675674 | 10920458 | 1243675287 |
| BlackFly40 | 8646086 | 1305558986 | 6611682 | 683202413 |
| BlackFly41 | 10029796 | 1514499196 | 7579750 | 873231109 |
| BlackFly42 | 14691830 | 2218466330 | 12196809 | 1443227901 |
| BlackFly43 | 14200262 | 2144239562 | 11087511 | 1274596527 |
| BlackFly44 | 12190272 | 1840731072 | 9998501 | 1147085995 |
| BlackFly45 | 12266724 | 1852275324 | 9756587 | 1106730398 |
| BlackFly46 | 10546382 | 1592503682 | 7436299 | 778570975 |
| BlackFly47 | 9580822 | 1446704122 | 7122148 | 799489096 |
| BlackFly48 | 11021364 | 1664225964 | 9059033 | 1015534398 |
| BlackFly49 | 9189200 | 1387569200 | 7021789 | 786538945 |
| BlackFly50 | 13864316 | 2093511716 | 12268048 | 1420847975 |
| BlackFly51 | 8788176 | 1327014576 | 6602351 | 690633816 |
| BlackFly52 | 12479582 | 1884416882 | 10616888 | 1243645802 |
| BlackFly53 | 16310048 | 2462817248 | 13489585 | 1583220590 |
| BlackFly55 | 8522448 | 1286889648 | 6572453 | 715669985 |
| BlackFly57 | 9687044 | 1462743644 | 7076582 | 760455742 |
| BlackFly58 | 9418760 | 1422232760 | 6903583 | 751600122 |
| BlackFly59 | 7709430 | 1164123930 | 5290944 | 541471964 |

Supplementary Table 2

| Accession | Family | Host | GenBank_Title |
| --- | --- | --- | --- |
| NC_078466.1 | Rhabdoviridae | Trifolium pratense | Red clover varicosavirus isolate HZ2 segment RNA2, complete sequence |
| NC_078467.1 | Rhabdoviridae | Trifolium pratense | Red clover varicosavirus isolate HZ2 segment RNA1, complete sequence |
| NC_079051.1 | Rhabdoviridae |  | MAG: Xinjiang varicosa-like virus isolate 237-k141_63393 segment 1, complete sequence |
| NC_079052.1 | Rhabdoviridae |  | MAG: Xinjiang varicosa-like virus isolate 237-k141_63393 segment 2, complete sequence |
| NC_079119.1 | Rhabdoviridae | Brassica rapa | Brassica rapa virus 1 N (QK880_s2gp1), protein 2 (QK880_s2gp2), and protein 3 (QK880_s2gp3) genes, complete cds |
| NC_079120.1 | Rhabdoviridae | Brassica rapa | Brassica rapa virus 1 L (QK880_s1gp1) gene, complete cds |
| NC_079121.1 | Rhabdoviridae | Lolium perenne | Lolium perenne virus 1 N (QK881_s2gp1), protein 2 (QK881_s2gp2), and protein 3 (QK881_s2gp3) genes, complete cds |
| NC_079122.1 | Rhabdoviridae | Lolium perenne | Lolium perenne virus 1 L (QK881_s1gp1) gene, complete cds |
| NC_079123.1 | Rhabdoviridae | Melampyrum roseum | Melampyrum roseum virus 1 L (QK882_s1gp1) gene, complete cds |
| NC_079124.1 | Rhabdoviridae | Melampyrum roseum | Melampyrum roseum virus 1 N (QK882_s2gp1), protein 2 (QK882_s2gp2), protein 3 (QK882_s2gp3), and protein 4 (QK882_s2gp4) genes, complete cds |
| NC_079125.1 | Rhabdoviridae | Zostera marina | Zostera associated varicosavirus 1 isolate Zoma segment RNA2 nucleocapsid (N), 40K protein 2 (P2), 34K protein 3 (P3), and 23K protein 4 (P4) genes, complete cds |
| NC_079126.1 | Rhabdoviridae | Zostera marina | Zostera associated varicosavirus 1 isolate Zoma segment RNA1 polymerase (L) gene, complete cds |
| NC_079127.1 | Rhabdoviridae | Allium angulosum | Allium angulosum virus 1 protein L (QK884_s1gp1) gene, complete cds |
| NC_079128.1 | Rhabdoviridae | Allium angulosum | Allium angulosum virus 1 protein N (QK884_s2gp1), protein 2 (QK884_s2gp2), and protein 3 (QK884_s2gp3) genes, complete cds |
| NC_075304.1 | Rhabdoviridae | Culex | Culex rhabdo-like virus strain CRVL/Los Angeles, complete genome |
| NC_075981.1 | Rhabdoviridae | Trichoprosopon theobaldi | Aruac virus nucleoprotein, phosphoprotein, matrix, glycoprotein, hypothetical proteins, and polymerase genes, complete cds |
| NC_075982.1 | Rhabdoviridae | Ixodes dentatus | Connecticut virus nucleoprotein, phosphoprotein, hypothetical protein, matrix, glycoprotein, hypothetical protein, and polymerase genes, complete cds |
| NC_075990.1 | Rhabdoviridae | Nyssomyia flaviscutellata | Inhangapi virus nucleoprotein, phosphoprotein, matrix, glycoprotein, hypothetical protein, and polymerase genes, complete cds |
| NC_076030.1 | Rhabdoviridae | Phlebotomus | Charleville virus strain Ch9824, partial genome |
| NC_076145.1 | Rhabdoviridae | Drosophila sturtevanti | Drosophila sturtevanti sigmavirus nucleocapsid protein (N), polymerase-associated protein (P), PP3 (X), matrix protein (M), glycoprotein (G), and RNA-dependent RNA polymerase (L) genes, complete cds |
| NC_076146.1 | Rhabdoviridae | Ceratitis capitata | Ceratitis capitata sigmavirus nucleocapsid protein (N), polymerase-associated protein (P), PP3 (X), matrix protein (M), glycoprotein (G), and RNA-dependent RNA polymerase (L) genes, complete cds |
| NC_076157.1 | Rhabdoviridae | Nematoda | Shayang ascaridia galli virus 2 strain HC21241 hypothetical protein 1 (QKL27_gp1), hypothetical protein 2 (QKL27_gp2), hypothetical protein 3 (QKL27_gp3), hypothetical protein 4 (QKL27_gp4), putative glycoprotein (QKL27_gp5), and RNA-dependent RNA polymerase (QKL27_gp6) genes, complete cds |
| NC_076163.1 | Rhabdoviridae | Solanum tuberosum | Potato yellow dwarf nucleorhabdovirus strain CYDV-constricta, complete genome |
| NC_076182.1 | Rhabdoviridae | Culex pseudovishnui | Culex pseudovishnui rhabdo-like virus 17NGK-Cps2-874 genes for nucleoprotein, phosphoprotein, hyppothetical protein, glycoprotein, RNA-dependent RNA polymerase, complete cds |
| NC_076208.1 | Rhabdoviridae | Ixodes scapularis | New Kent County virus isolate RTS126, complete genome |
| NC_076211.1 | Rhabdoviridae | Culex tarsalis | Dillard's Draw virus isolate DDrV-2015, complete genome |
| NC_076239.1 | Rhabdoviridae | Fragaria | Cytorhabdovirus fragariarugosus isolate A, complete genome |
| NC_076244.1 | Rhabdoviridae | Zanthoxylum | Green Sichuan pepper nucleorhabdovirus isolate ZPNu1, complete genome |
| NC_076250.1 | Polycipiviridae | Lasius neglectus | Lasius neglectus virus 2, complete genome |
| NC_076260.1 | Rhabdoviridae | Mansonia uniformis | Puchong virus isolate P5-350 Malaysian, complete genome |
| NC_076261.1 | Rhabdoviridae | Bos indicus | Hayes Yard virus isolate DPP4816, complete genome |
| NC_076267.1 | Rhabdoviridae | Malus domestica | Apple rootstock virus A, complete genome |
| NC_076289.1 | Rhabdoviridae | Trifolium pratense | Trifolium pratense virus B isolate 1/2014 putative N protein (QKM62_gp1), putative P protein (QKM62_gp2), putative P3 protein (QKM62_gp3), putative M protein (QKM62_gp4), putative G protein (QKM62_gp5), and putative L protein (QKM62_gp6) genes, complete cds |
| NC_076290.1 | Rhabdoviridae | Trifolium pratense | Trifolium pratense virus A isolate 29/15/1 putative N protein (QKM63_gp1), putative P protein (QKM63_gp2), putative P3 protein (QKM63_gp3), putative M protein (QKM63_gp4), putative G protein (QKM63_gp5), and putative L protein (QKM63_gp6) genes, complete cds |
| NC_076400.1 | Rhabdoviridae | Psorophora albigenu | UNVERIFIED: Lobeira virus isolate BR/MT_M05 polyprotein-like gene, partial sequence |
| NC_076415.1 | Rhabdoviridae | Amblyomma ovale | Blanchseco virus isolate TTP-Pool-17, complete genome |
| NC_076430.1 | Rhabdoviridae | Elettaria cardamomum | Cardamom vein clearing nucleorhabdovirus 1, complete genome |
| NC_076448.1 | Rhabdoviridae | Prunus persica | Peach virus 1 isolate NSTT, complete genome |
| NC_076472.1 | Rhabdoviridae | Ilex paraguariensis | Yerba mate virus A isolate Gob. Virasoro, complete genome |
| NC_076487.1 | Rhabdoviridae | Chlorion hirtum | Hymenopteran rhabdo-related virus OKIAV109 genomic sequence |
| NC_076489.1 | Rhabdoviridae | Pompilus cinereus | Hymenopteran rhabdo-related virus OKIAV38 genomic sequence |
| NC_076490.1 | Rhabdoviridae | Triodia sylvina | Lepidopteran rhabdo-related virus OKIAV34 genomic sequence |
| NC_076494.1 | Rhabdoviridae | Dipseliopoda | Bughendera virus isolate BF402 nucleoprotein (QKO67_gp1), phosphoprotein (QKO67_gp2), matrix protein (QKO67_gp3), glycoprotein (QKO67_gp4), and large protein (QKO67_gp5) genes, complete cds |
| NC_076501.1 | Rhabdoviridae | Heterodontonyx | Hymenopteran rhabdo-related virus OKIAV24 nucleoprotein (QKO74_gp1), hypothetical protein (QKO74_gp2), hypothetical protein (QKO74_gp3), glycoprotein (QKO74_gp4), and RdRp (QKO74_gp5) genes, complete cds |
| NC_076510.1 | Rhabdoviridae | Sonchus oleraceus | Sowthistle yellow vein virus isolate HWY65, complete genome |
| NC_076531.1 | Rhabdoviridae | Bacopa monnieri | Bacopa monnieri virus 1 isolate India, complete genome |
| NC_076532.1 | Rhabdoviridae | Bacopa monnieri | Bacopa monnieri virus 2 isolate India, complete genome |
| NC_076534.1 | Rhabdoviridae |  | Anole lyssa-like virus 1 A.allogus/Cuba/2011 RNA, complete genome |
| NC_076686.1 | Rhabdoviridae | Dryophytes cinereus | Frog lyssa-like virus 1 strain FLLV1-MaleA, complete genome |
| NC_076841.1 | Rhabdoviridae | Rhinolophus sinicus | Taiyi bat virus isolate 958, complete genome |
| NC_076842.1 | Rhabdoviridae | Rhinolophus sinicus | Yinshui bat virus isolate 1017, complete genome |
| NC_076864.1 | Rhabdoviridae |  | Paper mulberry mosaic-associated virus isolate SWU, complete genome |
| NC_076909.1 | Rhabdoviridae | Rosa hybrid cultivar | Rose virus R isolate MDR92016, complete genome |
| NC_076913.1 | Rhabdoviridae | Solanum aculeatissimum | Joa yellow blotch virus isolate Manaus, complete genome |
| NC_076914.1 | Rhabdoviridae | Chrysanthemum x morifolium | Chrysanthemum yellow dwarf virus isolate cq, complete genome |
| NC_076926.1 | Rhabdoviridae | Chrysura austriaca | Hymenopteran rhabdo-related virus isolate OKIAV23, complete sequence |
| NC_076927.1 | Rhabdoviridae | Chrysura radians | Hymenopteran rhabdo-related virus isolate OKIAV46, complete sequence |
| NC_076929.1 | Rhabdoviridae | Mansonia uniformis | Porton's virus isolate 0416MAL, partial genome |
| NC_076930.1 | Rhabdoviridae | Culex perfuscus | Bangoran virus isolate 0424RCA, partial genome |
| NC_076931.1 | Rhabdoviridae | Coquillettidia maculipennis | Boteke virus isolate 0417RCA, complete genome |
| NC_076932.1 | Rhabdoviridae |  | Sandjimba virus isolate 0408RCA, complete genome |
| NC_076933.1 | Rhabdoviridae | Eurillas virens | Nasoule virus isolate 0410RCA, complete genome |
| NC_076934.1 | Rhabdoviridae |  | Bimbo virus isolate 9716RCA, complete genome |
| NC_076935.1 | Rhabdoviridae |  | Kolongo virus isolate 9717RCA, complete genome |
| NC_076936.1 | Rhabdoviridae | Ploceus melanocephalus | Ouango virus isolate 9718RCA, complete genome |
| NC_076937.1 | Rhabdoviridae | Curruca curruca | Burg el Arab virus isolate 09023EGY, complete genome |
| NC_076938.1 | Rhabdoviridae | Curruca curruca | Matariya virus isolate 09027EGY, complete genome |
| NC_076939.1 | Rhabdoviridae | Rhinolophus ferrumequinum | Mediterranean bat virus isolate A09181, complete genome |
| NC_076970.1 | Rhabdoviridae | Agave tequilana | Agave tequilana virus 1 N (QKT51_gp1), P (QKT51_gp2), P3 (QKT51_gp3), M (QKT51_gp4), G (QKT51_gp5), and L (QKT51_gp6) genes, complete cds |
| NC_076971.1 | Rhabdoviridae | Asclepias syriaca | Asclepias syriaca virus 1 N (QKT52_gp1), P (QKT52_gp2), P3 (QKT52_gp3), M (QKT52_gp4), G (QKT52_gp5), P6 (QKT52_gp6), and L (QKT52_gp7) genes, complete cds |
| NC_076972.1 | Rhabdoviridae | Asclepias syriaca | Asclepias syriaca virus 2 N (QKT53_gp1), P (QKT53_gp2), P3 (QKT53_gp3), M (QKT53_gp4), G (QKT53_gp5), and L (QKT53_gp6) genes, complete cds |
| NC_076973.1 | Rhabdoviridae | Plectranthus aromaticus | Plectranthus aromaticus virus 1 N (QKT54_gp1), P (QKT54_gp2), P3 (QKT54_gp3), M (QKT54_gp4), G (QKT54_gp5), and L (QKT54_gp6) genes, complete cds |
| NC_076974.1 | Rhabdoviridae | Rhododendron delavayi | Rhododendron delavayi virus 1 N (QKT55_gp1), P (QKT55_gp2), P3 (QKT55_gp3), M (QKT55_gp4), G (QKT55_gp5), and L (QKT55_gp6) genes, complete cds |
| NC_076975.1 | Rhabdoviridae | Anthurium amnicola | Anthurium amnicola virus 1 N (QKT56_gp1), P (QKT56_gp2), P3 (QKT56_gp3), M (QKT56_gp4), G (QKT56_gp5), and L (QKT56_gp6) genes, complete cds |
| NC_076976.1 | Rhabdoviridae | Bemisia tabaci | Bemisia tabaci associated virus 1 N (QKT57_gp1), P (QKT57_gp2), P3 (QKT57_gp3), M (QKT57_gp4), G (QKT57_gp5), and L (QKT57_gp6) genes, complete cds |
| NC_076977.1 | Rhabdoviridae | Glehnia littoralis | Glehnia littoralis virus 1 N (QKT58_gp1), P (QKT58_gp2), P3 (QKT58_gp3), M (QKT58_gp4), G (QKT58_gp5), P6 (QKT58_gp6), and L (QKT58_gp7) genes, complete cds |
| NC_076978.1 | Rhabdoviridae | Gymnadenia x densiflora | Gymnadenia x densiflora virus 1 N (QKT59_gp1), P (QKT59_gp2), M (QKT59_gp3), and L (QKT59_gp4) genes, complete cds |
| NC_076979.1 | Rhabdoviridae | Nymphaea alba | Nymphaea alba virus 1 N (QKT60_gp1), P (QKT60_gp2), P3 (QKT60_gp3), M (QKT60_gp4), G (QKT60_gp5), P6 (QKT60_gp6), and L (QKT60_gp7) genes, complete cds |
| NC_076980.1 | Rhabdoviridae | Tagetes erecta | Tagetes erecta virus 1 N (QKT61_gp1), P (QKT61_gp2), P3 (QKT61_gp3), M (QKT61_gp4), and L (QKT61_gp5) genes, complete cds |
| NC_076981.1 | Rhabdoviridae | Trachyspermum ammi | Trachyspermum ammi virus 1 N (QKT62_gp1), P (QKT62_gp2), P3 (QKT62_gp3), M (QKT62_gp4), and L (QKT62_gp5) genes, complete cds |
| NC_076982.1 | Rhabdoviridae | Pinus flexilis | Pinus flexilis virus 1 N (QKT63_gp1), protein 2 (QKT63_gp2), protein 3 (QKT63_gp3), protein 4 (QKT63_gp4), and L (QKT63_gp5) genes, complete cds |
| NC_077111.1 | Rhabdoviridae | Trichobius sp. | Mejal virus isolate JAL10 nucleoprotein (N), phosphoprotein (P), matrix protein (M), glycoprotein (G), and RNA-dependent RNA polymerase (L) genes, complete cds |
| NC_077114.1 | Rhabdoviridae | Aves | Rhabdoviridae sp. isolate YSN900 genomic sequence |
| NC_077123.1 | Rhabdoviridae | Frankliniella intonsa | MAG: Hangzhou frankliniella intonsa rhabdovirus 1 isolate JM1FY86115, complete genome |
| NC_077129.1 | Rhabdoviridae | Rhinolophus pearsonii | Wufeng Rhinolophus pearsonii tupavirus 1, complete genome |
| NC_077130.1 | Rhabdoviridae | Leopoldamys edwardsi | Longquan Niviventer coninga ledantevirus 1, complete genome |
| NC_077151.1 | Rhabdoviridae | Cnidium officinale | Cnidium virus 1 isolate SK, complete genome |
| NC_077158.1 | Rhabdoviridae | Insecta | MAG: Xiangshan rhabdo-like virus 1 isolate Novel_23 nucleocapsid protein, hypothetical protein, spike glycoprotein, and RNA dependent RNA polymerase genes, complete cds |
| NC_077192.1 | Rhabdoviridae | Homo sapiens | Mundri virus isolate A14, partial genome |
| NC_055137.1 | Rhabdoviridae | Lepeophtheirus salmonis | Lepeophtheirus salmonis rhabdovirus No9, partial genome |
| NC_055138.1 | Rhabdoviridae | Lepeophtheirus salmonis | Lepeophtheirus salmonis rhabdovirus No127, partial genome |
| NC_055208.1 | Rhabdoviridae | Citrus sinensis | Citrus chlorotic spot virus strain Trs1 segment RNA1, complete sequence |
| NC_055290.1 | Rhabdoviridae | Culex sitiens | North Creek virus phosphoprotein (P) gene, partial cds |
| NC_055291.1 | Rhabdoviridae | Culex sitiens | North Creek virus glycoprotein (G) gene, complete cds |
| NC_055292.1 | Rhabdoviridae | Culex sitiens | North Creek virus RNA dependent RNA polymerase (L) gene, complete cds |
| NC_055293.1 | Rhabdoviridae | Culex sitiens | North Creek virus nucleoprotein (N) gene, partial cds |
| NC_055454.1 | Rhabdoviridae | Zea mays | Maize yellow striate virus, complete genome |
| NC_055456.1 | Rhabdoviridae | Culex | Kwatta virus nucleoprotein, phosphoprotein, hypothetical protein, matrix, hypothetical protein, glycoprotein, and polymerase genes, complete cds |
| NC_055457.1 | Rhabdoviridae | Haemaphysalis leporispalustris | New Minto virus nucleoprotein, phosphoprotein, matrix, glycoprotein, and polymerase genes, complete cds |
| NC_055460.1 | Rhabdoviridae | Culex annulirostris | Harrison Dam virus isolate CS75, partial genome |
| NC_055461.1 | Rhabdoviridae | Dermacentor variabilis | Sawgrass virus nucleoprotein, phosphoprotein, matrix, glycoprotein, and polymerase genes, complete cds |
| NC_055466.1 | Rhabdoviridae | Physostegia | Physostegia chlorotic mottle virus isolate PV-1182, complete genome |
| NC_055473.1 | Rhabdoviridae | Culex annulirostris | Holmes Jungle virus isolate DPP1163, complete genome |
| NC_055474.1 | Rhabdoviridae | Pipistrellus abramus | Taiwan bat lyssavirus isolate TWBLV/TN/2016, complete genome |
| NC_055477.1 | Rhabdoviridae | Ochlerotatus cantans | Ohlsdorf virus strain Germany/2012/Oc.cantans, complete genome |
| NC_055479.1 | Rhabdoviridae | Brassica oleracea | Cabbage cytorhabdovirus 1 strain FERA_050726, complete genome |
| NC_055484.1 | Rhabdoviridae | Triticum aestivum | Wheat yellow striate virus isolate SX-HC nucleocapsid protein, putative phosphoprotein, P3 protein, matrix protein, glycoprotein, hypothetical protein P6, and probable RNA-dependent RNA polymerase genes, complete cds |
| NC_055504.1 | Rhabdoviridae | Carica papaya | Papaya cytorhabdovirus isolate Los Rios_Ec, complete genome |
| NC_055505.1 | Rhabdoviridae | Ilex paraguariensis | Yerba mate chlorosis-associated virus isolate Montecarlo, complete genome |
| NC_055509.1 | Rhabdoviridae | Rhinella marina | Cuiaba virus strain BeAn 227841, partial genome |
| NC_055512.1 | Rhabdoviridae | Zea mays | Morogoro maize-associated virus isolate 16-0112 nucleocapsid protein, phosphoprotein, putative movement protein, matrix protein, glycoprotein, and RNA-dependent RNA polymerase genes, complete cds |
| NC_055529.1 | Rhabdoviridae | Rubus idaeus | Raspberry vein chlorosis virus isolate Hutton_1, complete genome |
| NC_055530.1 | Rhabdoviridae | Corythornis cristatus | Garba virus nucleoprotein, phosphoprotein, hypothetical protein, matrix, hypothetical protein, glycoprotein, and polymerase genes, complete cds |
| NC_055531.1 | Rhabdoviridae | Ochlerotatus sollicitans | Bahia Grande virus nucleoprotein, phosphoprotein, matrix, glycoprotein, hypothetical protein, and polymerase genes, complete cds |
| NC_055532.1 | Rhabdoviridae | Aedes sp. | Muir Springs virus nucleoprotein, phosphoprotein, matrix, glycoprotein, and polymerase genes, complete cds |
| NC_055567.1 | Rhabdoviridae | Fragaria x ananassa | Strawberry cytorhabdovirus 1 isolate B, complete genome |
| NC_052231.1 | Rhabdoviridae | Citrus sinensis | Citrus leprosis virus N strain ibi1 segment RNA2, complete sequence |
| NC_043065.1 | Rhabdoviridae | Drosophila tristis | Drosophila tristis sigmavirus RNA-dependent RNA polymerase (L) gene, partial cds |
| NC_043066.1 | Rhabdoviridae | Muscina stabulans | Muscina stabulans sigma virus RNA-dependent RNA polymerase (L) gene, partial cds |
| NC_043067.1 | Rhabdoviridae | Homo sapiens | Bas-Congo virus isolate BASV-1 N protein gene, partial cds |
| NC_043525.1 | Rhabdoviridae | Caligus rogercresseyi | Caligus rogercresseyi rhabdovirus strain CrRV-Ch01, partial genome |
| NC_043538.1 | Rhabdoviridae | Pipistrellus kuhlii | Vaprio virus nucleoprotein, phosphoprotein, matrix, glycoprotein, transcriptional unit 1, and polymerase genes, complete cds |
| NC_043649.1 | Rhabdoviridae | Clerodendrum sp. | Clerodendrum chlorotic spot virus isolate Prb1 segment RNA2, complete sequence |
| NC_040599.1 | Rhabdoviridae | Culex quinquefasciatus | Merida virus isolate MERD-Mex07, complete genome |
| NC_040602.1 | Rhabdoviridae | Aedes albopictus | Menghai rhabdovirus isolate Menghai, complete genome |
| NC_040664.1 | Rhabdoviridae | Hyalomma anatolicum anatolicum | Zahedan rhabdovirus isolate Ar Teh 157764, complete genome |
| NC_040669.1 | Rhabdoviridae | Ochlerotatus sp. | Riverside virus 1 strain RISV-Drava 1, complete genome |
| NC_040786.1 | Rhabdoviridae | Oryza sativa | Rice stripe mosaic virus isolate GD-LD, complete genome |
| NC_038236.1 | Rhabdoviridae | Equus caballus | Vesicular stomatitis Indiana virus strain 98COE, complete genome |
| NC_038275.1 | Rhabdoviridae | Culex sitiens | Mossuril virus nucleoprotein, phosphoprotein, hypothetical proteins, matrix, glycoprotein, hypothetical protein, and polymerase genes, complete cds |
| NC_038276.1 | Rhabdoviridae | Sus scrofa | Nishimuro virus viral cRNA for hypothetical proteins, complete cds |
| NC_038277.1 | Rhabdoviridae | Salmo trutta | Trout rhabdovirus 903/87 nucleocapsid protein, phosphoprotein, matrix protein, and glycoprotein genes, complete cds |
| NC_038278.1 | Rhabdoviridae | Drosophila affinis | Drosophila affinis sigmavirus nucleocapsid protein (N), polymerase-associated protein (P), PP3 (X), matrix protein (M), glycoprotein (G), and RNA-dependent RNA polymerase (L) genes, complete cds |
| NC_038279.1 | Rhabdoviridae | Drosophila ananassae | Drosophila ananassae sigmavirus nucleocapsid protein (N), polymerase-associated protein (P), PP3 (X), matrix protein (M), glycoprotein (G), and RNA-dependent RNA polymerase (L) genes, complete cds |
| NC_038280.1 | Rhabdoviridae | Diptera | Drosophila immigrans sigmavirus strain SCM45623 nucleocapsid protein, polymerase-associated protein, PP3, matrix protein, glycoprotein, and RNA-dependent RNA polymerase genes, complete cds |
| NC_038281.1 | Rhabdoviridae | Drosophila melanogaster | Drosophila melanogaster sigma virus HAP23, complete genome |
| NC_038282.1 | Rhabdoviridae | Homo sapiens | Ekpoma virus 1 isolate EKV-1, partial genome |
| NC_038283.1 | Rhabdoviridae | Homo sapiens | Ekpoma virus 2 isolate EKV-2, partial genome |
| NC_038284.1 | Rhabdoviridae | Aves | Durham virus nucleocapsid, phosphoprotein, putative protein C, matrix protein, small hydrophobic protein, and glycoprotein genes, complete cds |
| NC_038285.1 | Rhabdoviridae | Lutzomyia | Carajas virus nucleoprotein, phosphoprotein, matrix, glycoprotein, and polymerase genes, complete cds |
| NC_038286.1 | Rhabdoviridae | Philander opossum | Piry virus strain BeAn2423, complete genome |
| NC_038287.1 | Rhabdoviridae | Phlebotomus perfiliewi | Radi virus nucleoprotein, phosphoprotein, matrix, glycoprotein, and polymerase genes, complete cds |
| NC_038755.1 | Rhabdoviridae |  | Coffee ringspot virus strain Lavras segment RNA2, complete sequence |
| NC_039020.1 | Rhabdoviridae | Dipseliopoda | Kanyawara virus isolate MPK004 nucleoprotein, phosphoprotein, matrix, glycoprotein, and polymerase genes, complete cds |
| NC_039021.1 | Rhabdoviridae | Culicoides peregrinus | Beatrice Hill virus isolate CSIRO 25, complete genome |
| NC_039200.1 | Rhabdoviridae | Psorophora albigenu | Balsa almendravirus, complete genome |
| NC_039201.1 | Rhabdoviridae | Culicoides | Curionopolis virus nucleoprotein, phosphoprotein, matrix, hypothetical proteins, glycoprotein, hypothetical proteins, and polymerase genes, complete cds |
| NC_039202.1 | Rhabdoviridae | Culiseta melanura | Flanders virus nucleoprotein, phosphoprotein, hypothetical proteins, matrix, glycoprotein, hypothetical protein, and polymerase genes, complete cds |
| NC_039206.1 | Rhabdoviridae | Haemagogus | Jurona virus nucleoprotein, phosphoprotein, matrix, glycoprotein, and polymerase genes, complete cds |
| NC_036390.1 | Rhabdoviridae | Zea mays | Maize Iranian mosaic nucleorhabdovirus, complete genome |
| NC_031957.1 | Rhabdoviridae | Anopheles quadrimaculatus | Coot Bay virus strain EVG5-53, complete sequence |
| NC_031958.1 | Rhabdoviridae | Culicidae | Rio Chico virus strain GAM 195, complete sequence |
| NC_035132.1 | Rhabdoviridae | Culex quinquefasciatus | Culex rhabdo-like virus strain mosWSB71420, complete genome |
| NC_034508.1 | Rhabdoviridae | Lutzomyia | Morreton virus nucleoprotein, phosphoprotein, matrix, glycoprotein, and polymerase genes, complete cds |
| NC_034529.1 | Rhabdoviridae | Ameiva ameiva | Sena Madureira virus nucleoprotein, hypothetical protein, phosphoprotein, matrix, glycoprotein, hypothetical protein, and polymerase genes, complete cds |
| NC_034530.1 | Rhabdoviridae | Ameiva ameiva | Marco virus nucleoprotein, phosphoprotein, matrix, glycoprotein, hypothetical proteins, and polymerase genes, complete cds |
| NC_034531.1 | Rhabdoviridae | Culex tarsalis | Manitoba virus nucleoprotein, phosphoprotein, hypothetical proteins, matrix, glycoprotein, hypothetical protein, and polymerase genes, complete cds |
| NC_034533.1 | Rhabdoviridae | Riparia paludicola | Landjia virus nucleoprotein, phosphoprotein, hypothetical proteins, matrix, glycoprotein, hypothetical proteins, and polymerase genes, complete cds |
| NC_034534.1 | Rhabdoviridae | Coquillettidia albicosta | Rochambeau virus nucleoprotein, phosphoprotein, matrix, hypothetical proteins, glycoprotein, hypothetical proteins, and polymerase genes, complete cds |
| NC_034535.1 | Rhabdoviridae | Rattus rattus | Barur virus nucleoprotein, phosphoprotein, matrix, glycoprotein, and polymerase genes, complete cds |
| NC_034536.1 | Rhabdoviridae | Culicoides | Itacaiunas virus nucleoprotein, phosphoprotein, matrix, hypothetical protein, glycoprotein, hypothetical protein, and polymerase genes, complete cds |
| NC_034537.1 | Rhabdoviridae | Culex dunni | La Joya virus nucleoprotein, phosphoprotein, hypothetical proteins, matrix, hypothetical proteins, glycoprotein, hypothetical proteins, and polymerase genes, complete cds |
| NC_034538.1 | Rhabdoviridae | Culicinae | Joinjakaka virus nucleoprotein, phosphoprotein, hypothetical protein, matrix, glycoprotein, hypothetical proteins, and polymerase genes, complete cds |
| NC_034539.1 | Rhabdoviridae | Eretmapodites leucopous | Nkolbisson virus nucleoprotein, phosphoprotein, matrix, glycoprotein, and polymerase genes, complete cds |
| NC_034540.1 | Rhabdoviridae | Gerbilliscus kempi | Keuraliba virus nucleoprotein, phosphoprotein, matrix, glycoprotein, hypothetical protein, and polymerase genes, complete cds |
| NC_034541.1 | Rhabdoviridae | Culex tarsalis | Gray Lodge virus nucleoprotein, phosphoprotein, hypothetical proteins, matrix, glycoprotein, hypothetical protein, and polymerase genes, complete cds |
| NC_034542.1 | Rhabdoviridae | Sergentomyia | Sripur virus nucleoprotein, hypothetical protein, phosphoprotein, hypothetical protein, matrix, hypothetical protein, glycoprotein, hypothetical protein, and polymerase genes, complete cds |
| NC_034543.1 | Rhabdoviridae | Culex annulirostris | Ord River virus nucleoprotein, hypothetical protein, phosphoprotein, hypothetical proteins, matrix, glycoprotein, hypothetical protein, and polymerase genes, complete cds |
| NC_034544.1 | Rhabdoviridae | Lutzomyia | Iriri virus nucleoprotein, phosphoprotein, matrix, hypothetical proteins, glycoprotein, hypothetical proteins, and polymerase genes, complete cds |
| NC_034545.1 | Rhabdoviridae | Rhinolophus eloquens | Mount Elgon bat virus nucleoprotein, phosphoprotein, matrix, glycoprotein, and polymerase genes, complete cds |
| NC_034546.1 | Rhabdoviridae | Culicoides insignis | Sweetwater Branch virus nucleoprotein, phosphoprotein, matrix, hypothetical proteins, glycoprotein, hypothetical protein, and polymerase genes, complete cds |
| NC_034548.1 | Rhabdoviridae | Rhinolophus cornutus | Oita virus nucleoprotein, phosphoprotein, matrix, glycoprotein, and polymerase genes, complete cds |
| NC_034549.1 | Rhabdoviridae | Microtus montanus | Klamath virus nucleoprotein, phosphoprotein, hypothetical protein, matrix, hypothetical protein, glycoprotein, hypothetical protein, and polymerase genes, complete cds |
| NC_034550.1 | Rhabdoviridae | Ameiva ameiva | Chaco virus nucleoprotein, hypothetical protein, phosphoprotein, matrix, hypothetical protein, glycoprotein, hypothetical protein, and polymerase genes, complete cds |
| NC_034551.1 | Rhabdoviridae | Colocasia esculenta | Colocasia bobone disease-associated virus strain SI, complete genome |
| NC_034443.1 | Rhabdoviridae | Homo sapiens | Le Dantec virus nucleoprotein, phosphoprotein, matrix, glycoprotein, hypothetical protein, and polymerase genes, complete cds |
| NC_034447.1 | Rhabdoviridae | Culex tarsalis | Hart Park virus nucleoprotein, phosphoprotein, hypothetical proteins, matrix, glycoprotein, hypothetical protein, and polymerase genes, complete cds |
| NC_034448.1 | Rhabdoviridae | Culex portesi | Mosqueiro virus nucleoprotein, phosphoprotein, hypothetical proteins, matrix, glycoprotein, hypothetical protein, and polymerase genes, complete cds |
| NC_034449.1 | Rhabdoviridae | Culex annulirostris | Parry Creek virus nucleoprotein, phosphoprotein, hypothetical proteins, matrix, glycoprotein, hypothetical protein, and polymerase genes, complete cds |
| NC_034450.1 | Rhabdoviridae | Culex annulioris | Kamese virus nucleoprotein, phosphoprotein, hypothetical proteins, matrix, glycoprotein, hypothetical protein, and polymerase genes, complete cds |
| NC_034451.1 | Rhabdoviridae | Myotis yumanensis | Kern Canyon virus nucleoprotein, phosphoprotein, matrix, glycoprotein, hypothetical protein, and polymerase genes, complete cds |
| NC_034454.1 | Rhabdoviridae | Culicoides punctatus | Fukuoka virus nucleoprotein, phosphoprotein, matrix, hypothetical protein, glycoprotein, and polymerase genes, complete cds |
| NC_034240.1 | Rhabdoviridae | Solanum lycopersicum | Tomato yellow mottle-associated virus, complete genome |
| NC_033701.1 | Rhabdoviridae | Nematoda | Xingshan nematode virus 4 strain XSNXC32924 putative nucleoprotein, hypothetical protein 1, hypothetical protein 2, putative glycoprotein, and RNA-dependent RNA polymerase genes, complete cds |
| NC_033705.1 | Rhabdoviridae | Nematoda | Xinzhou nematode virus 4 strain XZSJSC65771 putative nucleoprotein, hypothetical protein 2, hypothetical protein 3, putative glycoprotein, and RNA-dependent RNA polymerase genes, complete cds |
| NC_032739.1 | Rhabdoviridae | Crustacea | Wenling crustacean virus 10 strain WLJQ101844 hypothetical protein 1, hypothetical protein 2, hypothetical protein 3, putative glycoprotein, and RNA-dependent RNA polymerase genes, complete cds |
| NC_032781.1 | Rhabdoviridae | Crustacea | Wenling crustacean virus 11 strain WLJQ201798 hypothetical protein 1, hypothetical protein 2, hypothetical protein 3, putative glycoprotein, and RNA-dependent RNA polymerase genes, complete cds |
| NC_032907.1 | Rhabdoviridae | Lepidoptera | Hubei lepidoptera virus 2 strain LCM101902 putative nucleoprotein, hypothetical protein 2, putative membrane protein, putative glycoprotein 1, putative glycoprotein 2, and RNA-dependent RNA polymerase genes, complete cds |
| NC_033034.1 | Rhabdoviridae | Diptera | Hubei diptera virus 9 strain SCM172232 putative nucleoprotein, hypothetical protein 2, putative X protein, putative matrix protein, putative glycoprotein, and RNA-dependent RNA polymerase genes, complete cds |
| NC_033070.1 | Rhabdoviridae | Diptera | Hubei dimarhabdovirus virus 1 strain SCM51525 putative nucleoprotein, hypothetical protein, putative matrix protein, putative glycoprotein, and RNA-dependent RNA polymerase genes, complete cds |
| NC_033103.1 | Rhabdoviridae | Diptera | Hubei diptera virus 10 strain SCM43656 putative nucleoprotein, hypothetical protein, putative X protein, putative matrix protein, putative glycoprotein, and RNA-dependent RNA polymerase genes, complete cds |
| NC_033267.1 | Rhabdoviridae | Ascaris suum | Hubei rhabdo-like virus 9 strain WHZHC73015 hypothetical protein 1, hypothetical protein 2, hypothetical protein 3, hypothetical protein 4, hypothetical protein 5, putative glycoprotein, and RNA-dependent RNA polymerase genes, complete cds |
| NC_031988.1 | Rhabdoviridae | Pteropus giganteus | Gannoruwa bat lyssavirus isolate RV3266, complete genome |
| NC_031955.1 | Rhabdoviridae | Miniopterus schreibersii | Lleida bat lyssavirus isolate RV3208, complete genome |
| NC_031301.1 | Rhabdoviridae | Hippoboscidae | Wuhan Louse Fly Virus 5 strain BFJSC-5 nucleocapsid (N), phosphoprotein (P), matrix protein (M), glycoprotein (G), and RNA-dependent RNA polymerase (L) genes, complete cds |
| NC_031302.1 | Rhabdoviridae | Hippoboscidae | Wuhan Louse Fly Virus 9 strain BFJSC-7 nucleocapsid (N), putative phosphoprotein (ORF2), matrix protein (M), glycoprotein (G), and RNA-dependent RNA polymerase (L) genes, complete cds |
| NC_031304.1 | Rhabdoviridae | Rhipicephalus microplus | Wuhan Tick Virus 1 strain X78-2 nucleocapsid (N), ORF2 (ORF2), ORF3 (ORF3), and RNA-dependent RNA polymerase (L) genes, complete cds |
| NC_031305.1 | Rhabdoviridae | Haemaphysalis hystricis | Yongjia Tick Virus 2 strain YJ1-2 nucleocapsid (N), putative phosphoprotein (ORF2), matrix protein (M), glycoprotein (G), and RNA-dependent RNA polymerase (L) genes, complete cds |
| NC_031225.1 | Rhabdoviridae | Hyalopterus pruni | Wuhan Insect virus 4 strain YCYC03 nucleocapsid (N), phosphoprotein (P), 4b protein (4b), putative matrix protein (ORF4), glycoprotein (G), and RNA-dependent RNA polymerase (L) genes, complete cds |
| NC_031227.1 | Rhabdoviridae | Hyalopterus pruni | Wuhan Insect virus 5 strain YCYC02 nucleocapsid (N), phosphoprotein (P), 4b protein (4b), matrix protein (M), glycoprotein (G), and RNA-dependent RNA polymerase (L) genes, complete cds |
| NC_031232.1 | Rhabdoviridae | Hyalopterus pruni | Wuhan Insect virus 6 strain SXCC01-1 nucleocapsid (N), phosphoprotein (P), 4b protein (4b), matrix protein (M), glycoprotein (G), and RNA-dependent RNA polymerase (L) genes, complete cds |
| NC_031240.1 | Rhabdoviridae | Hippoboscidae | Wuhan Louse Fly Virus 10 strain BFJSC-8 nucleocapsid (N), putative phosphoprotein (ORF2), matrix protein (M), glycoprotein (G), and RNA-dependent RNA polymerase (L) genes, complete cds |
| NC_031278.1 | Rhabdoviridae | Musca domestica | Wuhan Fly Virus 2 strain SYY1-3 nucleocapsid (N), ORF2 (ORF2), X protein (X), matrix protein (M), glycoprotein (G), and RNA-dependent RNA polymerase (L) genes, complete cds |
| NC_031282.1 | Rhabdoviridae | Musca domestica | Wuhan House Fly Virus 1 strain SYY2-4 nucleocapsid (N), putative phosphoprotein (ORF2), putative X protein (ORF3), matrix protein (M), glycoprotein (G), and RNA-dependent RNA polymerase (L) genes, complete cds |
| NC_031283.1 | Rhabdoviridae | Musca domestica | Wuhan House Fly Virus 2 strain SYY4-5 ORF1 (ORF1), ORF2 (ORF2), ORF3 (ORF3), glycoprotein (G), and RNA-dependent RNA polymerase (L) genes, complete cds |
| NC_031215.1 | Rhabdoviridae | Musca domestica | Shayang Fly Virus 2 strain SYY1-8 nucleocapsid (N), ORF2 (ORF2), X protein (X), matrix protein (M), glycoprotein (G), and RNA-dependent RNA polymerase (L) genes, complete cds |
| NC_031216.1 | Rhabdoviridae | Chrysomya megacephala | Shayang Fly Virus 3 strain SYY1-1 ORF1 (ORF1), ORF2 (ORF2), ORF3 (ORF3), glycoprotein (G), and RNA-dependent RNA polymerase (L) genes, complete cds |
| NC_031079.1 | Rhabdoviridae | Hyalomma asiaticum | Bole Tick Virus 2 strain BL076 nucleocapsid (N), putative phosphoprotein (ORF2), matrix protein (M), glycoprotein (G), and RNA-dependent RNA polymerase (L) genes, complete cds |
| NC_028484.1 | Rhabdoviridae | Culex bitaeniorhynchus | Tongilchon virus 1 strain A12.2676/ROK/2012, complete genome |
| NC_028255.1 | Rhabdoviridae |  | Cocal virus Indiana 2, complete genome |
| NC_028246.1 | Rhabdoviridae | Bos taurus | Adelaide River virus isolate DPP61, complete genome |
| NC_028231.1 | Rhabdoviridae | Thunbergia alata | Datura yellow vein virus, complete genome |
| NC_028232.1 | Rhabdoviridae | Culicoides austropalpalis | Walkabout Creek virus isolate CS1056, complete genome |
| NC_028234.1 | Rhabdoviridae | Psychodidae | Santa barbara virus strain AR775619, complete genome |
| NC_028236.1 | Rhabdoviridae | Eidolon helvum | Kumasi rhabdbovirus, complete genome |
| NC_028237.2 | Rhabdoviridae | Medicago sativa | Alfalfa dwarf virus isolate Manfredi, complete genome |
| NC_028239.1 | Rhabdoviridae | Bos taurus | Koolpinyah virus isolate DPP819, complete genome |
| NC_028241.1 | Rhabdoviridae | Mansonia uniformis | Yata virus isolate DakArB 2181, complete genome |
| NC_028244.1 | Rhabdoviridae |  | Barley yellow striate mosaic virus strain Hebei, complete genome |
| NC_026798.1 | Rhabdoviridae |  | Black grass varicosavirus-like virus segment RNA 2 |
| NC_026801.1 | Rhabdoviridae |  | Black grass varicosavirus-like virus segment RNA 1 |
| NC_025385.1 | Rhabdoviridae |  | Khujand lyssavirus, complete genome |
| NC_025387.1 | Rhabdoviridae | Scophthalmus maximus | Scophthalmus maximus rhabdovirus, complete genome |
| NC_025389.1 | Rhabdoviridae | Agapanthus | Eggplant mottled dwarf virus isolate Agapanthus, complete genome |
| NC_025391.1 | Rhabdoviridae | Cryptoblepharus virgatus | Almpiwar virus isolate MRM4059, complete genome |
| NC_025393.1 | Rhabdoviridae | Culicidae | Arboretum virus isolate Lo-121, complete genome |
| NC_025394.1 | Rhabdoviridae | Culicidae | Perinet virus, complete genome |
| NC_025395.1 | Rhabdoviridae | Ochlerotatus fulvus | Puerto Almendras virus isolate LO-39, complete genome |
| NC_025396.1 | Rhabdoviridae | Bos taurus | Kimberley virus isolate CS368, complete genome |
| NC_025397.1 | Rhabdoviridae | Bos taurus | Coastal Plains virus strain DPP53, complete genome |
| NC_025399.1 | Rhabdoviridae | Culex edwardsi | Oak-Vale virus strain CSIRO 1342, complete genome |
| NC_025400.1 | Rhabdoviridae | Mansonia uniformis | Malakal virus isolate SudAr 1169-64, complete genome |
| NC_025401.1 | Rhabdoviridae | Gallus gallus | Sunguru virus isolate Ug#41, complete genome |
| NC_025405.1 | Rhabdoviridae |  | Niakha virus isolate DakArD 88909, complete genome |
| NC_025406.1 | Rhabdoviridae | Lagenorhynchus albirostris | Dolphin rhabdovirus isolate pxV1 1992, complete genome |
| NC_025382.1 | Rhabdoviridae | Spodoptera frugiperda | Spodoptera frugiperda rhabdovirus isolate Sf, complete genome |
| NC_025384.1 | Rhabdoviridae | Culex tritaeniorhynchus | Culex tritaeniorhynchus rhabdovirus RNA, complete genome, strain: TY |
| NC_025340.1 | Rhabdoviridae | Amblyomma americanum | Long Island tick rhabdovirus strain LS1, complete genome |
| NC_025341.1 | Rhabdoviridae | Macronycteris commersoni | Fikirini bat rhabdovirus isolate KEN352, complete genome |
| NC_025342.1 | Rhabdoviridae | Amblyomma | Kolente virus isolate DakAr K7292, complete genome |
| NC_025353.1 | Rhabdoviridae | Equus asinus x Equus caballus | Vesicular stomatitis Alagoas virus Indiana 3, complete genome |
| NC_025356.1 | Rhabdoviridae | Esox lucius | Pike fry rhabdovirus isolate F4, complete genome |
| NC_025358.1 | Rhabdoviridae | Bos taurus | Berrimah virus strain DPP 63, complete genome |
| NC_025359.1 | Rhabdoviridae | Culex decens | Moussa virus isolate C23, complete genome |
| NC_025362.1 | Rhabdoviridae | Sabethes intermedius | Xiburema virus isolate XIBV/BE AR 362159, complete genome |
| NC_025364.1 | Rhabdoviridae | Ochlerotatus campestris | Malpais Spring virus strain 85-488NM, complete genome |
| NC_025365.1 | Rhabdoviridae | Macronycteris commersoni | Shimoni bat virus, complete genome |
| NC_025371.1 | Rhabdoviridae | Tinca tinca | Tench rhabdovirus S64, complete genome |
| NC_025376.1 | Rhabdoviridae | Ctenopharyngodon idella | Grass carp rhabdovirus V76, complete genome |
| NC_025377.1 | Rhabdoviridae |  | West Caucasian bat virus, complete genome |
| NC_025378.1 | Rhabdoviridae |  | Yug Bogdanovac virus, complete genome |
| NC_025251.1 | Rhabdoviridae | Myotis nattereri | Bokeloh bat lyssavirus isolate 21961, complete genome |
| NC_025255.1 | Rhabdoviridae |  | Maraba virus from Brazil, complete genome |
| NC_024473.1 | Rhabdoviridae | Bos taurus | Vesicular stomatitis New Jersey virus isolate NJ1184HDB, complete genome |
| NC_022755.1 | Rhabdoviridae | Eptesicus fuscus | American bat vesiculovirus TFFN-2013 isolate liver2008, complete genome |
| NC_022580.1 | Rhabdoviridae | Drosophila obscura | Drosophila obscura sigma virus 10A, complete genome |
| NC_022581.1 | Rhabdoviridae | Anguilla anguilla | Eel Virus European X complete genome, viral cRNA, isolate 153311 |
| NC_020803.1 | Rhabdoviridae | Perca fluviatilis | Perch perhabdovirus isolate PRV nucleoprotein (N) gene, nucleocapsid (N) gene, complete cds |
| NC_020804.1 | Rhabdoviridae | Culicoides brevitarsis | Tibrogargan virus strain CS132, complete genome |
| NC_020805.1 | Rhabdoviridae | Homo sapiens | Chandipura virus isolate CIN 0451, complete genome |
| NC_020806.1 | Rhabdoviridae | Phlebotomus papatasi | Isfahan virus N gene, P gene, M gene, G gene and L gene, genomic RNA |
| NC_020807.1 | Rhabdoviridae | Eidolon helvum | Lagos bat virus isolate 0406SEN, complete genome |
| NC_020808.1 | Rhabdoviridae |  | Aravan virus, complete genome |
| NC_020809.1 | Rhabdoviridae |  | Irkut virus, complete genome |
| NC_020810.1 | Rhabdoviridae | Homo sapiens | Duvenhage virus isolate 86132SA, complete genome |
| NC_018629.1 | Rhabdoviridae | Civettictis civetta | Ikoma lyssavirus, complete genome |
| NC_018381.2 | Rhabdoviridae | Diospyros kaki | Persimmon virus A viral cRNA, complete genome, clone: Kaki13-14 |
| NC_017685.1 | Rhabdoviridae | Mansonia uniformis | Obodhiang virus, complete genome |
| NC_017714.1 | Rhabdoviridae | Culicoides | Kotonkan virus, complete genome |
| NC_016136.1 | Rhabdoviridae |  | Potato yellow dwarf virus, complete genome |
| NC_013955.1 | Rhabdoviridae |  | Ngaingan virus, complete genome |
| NC_011639.1 | Rhabdoviridae | Culicoides austropalpalis | Wongabel virus, complete genome |
| NC_011568.1 | Rhabdoviridae |  | Lettuce big-vein associated virus segment 2, complete genome |
| NC_011532.1 | Rhabdoviridae | Lactuca sativa | Lettuce yellow mottle virus, complete genome |
| NC_009609.1 | Rhabdoviridae |  | Orchid fleck virus genomic RNA, segment RNA 2, complete sequence |
| NC_009527.1 | Rhabdoviridae | Eptesicus serotinus | European bat lyssavirus 1, complete genome |
| NC_009528.2 | Rhabdoviridae | Homo sapiens | European bat lyssavirus 2 isolate RV1333, complete genome |
| NC_008514.1 | Rhabdoviridae | Siniperca chuatsi | Siniperca chuatsi rhabdovirus, complete genome |
| NC_007642.1 | Rhabdoviridae | Allium sativum | Lettuce necrotic yellows virus, complete genome |
| NC_007020.1 | Rhabdoviridae |  | Tupaia virus, complete genome |
| NC_006942.1 | Rhabdoviridae | Colocasia esculenta | Taro vein chlorosis virus, complete genome |
| NC_006429.1 | Rhabdoviridae |  | Mokola virus, complete genome |
| NC_005974.1 | Rhabdoviridae |  | Maize fine streak virus, complete genome |
| NC_005975.1 | Rhabdoviridae |  | Maize mosaic virus, complete genome |
| NC_005093.1 | Rhabdoviridae | Oncorhynchus mykiss | Hirame rhabdovirus, complete genome |
| NC_002526.1 | Rhabdoviridae |  | Bovine ephemeral fever virus, complete genome |
| NC_002251.1 | Rhabdoviridae |  | Northern cereal mosaic virus, complete genome |
| NC_000903.1 | Rhabdoviridae | Channa striata | Snakehead rhabdovirus complete genome |
| NC_003243.1 | Rhabdoviridae |  | Australian bat lyssavirus, complete genome |
| NC_000855.1 | Rhabdoviridae |  | Viral hemorrhagic septicemia virus Fil3, complete genome |
| NC_003746.1 | Rhabdoviridae |  | Rice yellow stunt virus, complete genome |
| NC_002803.1 | Rhabdoviridae | Cyprinus carpio | Spring viraemia of carp virus, complete genome |
| NC_001652.1 | Rhabdoviridae | Oncorhynchus tshawytscha | Infectious hematopoietic necrosis virus, complete genome |
| NC_001615.3 | Rhabdoviridae | Nicotiana x edwardsonii | Sonchus yellow net virus complete genome |
| NC_001542.1 | Rhabdoviridae |  | Rabies virus, complete genome |
| OZ077875.1 | Rhabdoviridae | Onchocerca volvulus | Onchocerca volvulus RNA Virus 1 isolate missing: third party data genome assembly, chromosome: OvRV1 |
| KP688058.1 | Flaviviridae | Culex | Mercadeo virus isolate ER-M10, complete genome |
| KT192549.1 | Flaviviridae |  | Parramatta River virus isolate 92-B115745, complete genome |
| AB981186.1 | Flaviviridae | Culex | Mosquito flavivirus gene for polyprotein, complete cds, strain: YDFV/Oct/2013 |
| KF917536.1 | Flaviviridae | Formicarius analis | Cacipacore virus strain BeAn 3276000, complete genome |
| KJ469370.1 | Flaviviridae | Cynopterus brachyotis | Batu Cave virus strain P70-1459, complete genome |
| KJ469371.1 | Flaviviridae | Sigmodon hispidus | Jutiapa virus strain JG-128, complete genome |
| KJ469372.1 | Flaviviridae | Cynopterus brachyotis | Phnom Penh bat virus strain 30834_A38, complete genome |
| KF815939.1 | Flaviviridae | Ixodes uriae | Tyuleniy virus strain LEIV-6C polyprotein gene, complete cds |
| KC464457.1 | Flaviviridae | Culex tritaeniorhynchus | Mosquito flavivirus isolate LSFlaviV-A20-09, complete genome |
| KC505248.1 | Flaviviridae | Coquillettidia xanthogaster | Palm Creek virus isolate 56 polyprotein gene, complete cds |
| JQ268258.1 | Flaviviridae | Culicidae | Hanko virus polyprotein gene, complete cds |
| JX236040.3 | Flaviviridae |  | Ntaya virus isolate IPDIA, complete genome |
| HE574574.1 | Flaviviridae | Culex theileri | Culex theileri flavivirus RP-2011 gene for viral polyprotein, genomic RNA, isolate 178 |
| JF895923.2 | Flaviviridae | Anatidae | Tembusu virus strain JS804, complete genome |
| DQ859056.1 | Flaviviridae |  | Banzi virus strain SAH 336 polyprotein gene, complete cds |
| DQ859057.1 | Flaviviridae |  | Bouboui virus strain DAK AR B490 polyprotein gene, complete cds |
| DQ859060.1 | Flaviviridae |  | Edge Hill virus strain YMP 48 polyprotein gene, complete cds |
| DQ859062.1 | Flaviviridae |  | Saboya virus strain Dak AR D4600 polyprotein gene, complete cds |
| DQ859065.1 | Flaviviridae |  | Uganda S virus polyprotein gene, complete cds |
| DQ859066.1 | Flaviviridae |  | Jugra virus strain P-9-314 polyprotein gene, complete cds |
| DQ859067.1 | Flaviviridae |  | Potiskum virus strain IBAN 10069 polyprotein gene, complete cds |
| GQ165809.2 | Flaviviridae | Mansonia africana | Nakiwogo virus strain Uganda08 polyprotein gene, partial cds |
| AB488408.1 | Flaviviridae | Aedes albopictus | Aedes flavivirus genomic RNA, complete genome, strain: Narita-21 |
| EU707555.1 | Flaviviridae |  | Wesselsbron virus strain SAH177, complete genome |
| FJ644291.1 | Flaviviridae | Culex tritaeniorhynchus | Quang Binh virus isolate VN180, complete genome |
| AB377213.1 | Flaviviridae | Culex pipiens | Culex flavivirus genomic RNA, complete genome, strain: NIID-21-2 |
| DQ837641.1 | Flaviviridae | Chiroptera | Entebbe bat virus strain UgIL-30, complete genome |
| DQ837642.1 | Flaviviridae | Culicidae | Sepik virus strain MK7148, complete genome |
| DQ525916.1 | Flaviviridae |  | St. Louis encephalitis virus strain Kern217, complete genome |
| DQ235144.1 | Flaviviridae |  | Meaban virus from France polyprotein gene, complete cds |
| DQ235145.1 | Flaviviridae |  | Gadgets Gully virus from Australia polyprotein gene, complete cds |
| DQ235146.1 | Flaviviridae |  | Kadam virus from Uganda polyprotein gene, complete cds |
| DQ235149.1 | Flaviviridae |  | Royal Farm virus from Afghanistan polyprotein gene, complete cds |
| DQ235150.1 | Flaviviridae |  | Saumarez Reef virus from Australia polyprotein gene, complete cds |
| DQ235151.1 | Flaviviridae |  | Turkish sheep encephalitis virus polyprotein gene, complete cds |
| DQ235152.1 | Flaviviridae |  | Spanish sheep encephalitis virus polyprotein gene, complete cds |
| DQ235153.1 | Flaviviridae |  | Greek goat encephalitis virus polyprotein gene, complete cds |
| AY323490.1 | Flaviviridae |  | Kyasanur forest disease virus polyprotein gene, complete cds |
| AY632535.2 | Flaviviridae | Simiiformes | Zika virus strain MR 766, complete genome |
| AY632536.4 | Flaviviridae |  | Bussuquara virus strain BeAn 4073, complete genome |
| AY632539.4 | Flaviviridae |  | Ilheus virus strain Original, complete genome |
| AY632540.2 | Flaviviridae | Culicidae | Kedougou virus strain DakAar D1470, complete genome |
| AY632541.4 | Flaviviridae |  | Kokobera virus strain AusMRM 32, complete genome |
| AY632545.2 | Flaviviridae | Culicidae | Bagaza virus strain DakAr B209, complete genome |
| AY453411.1 | Flaviviridae |  | Usutu virus strain Vienna 2001 from Austria, complete genome |
| AY193805.1 | Flaviviridae |  | Omsk hemorrhagic fever virus strain Bogoluvovska, complete genome |
| AB114858.1 | Flaviviridae | Chiroptera | Yokose virus genomic RNA, complete genome, strain:Oita 36 |
| AY149905.1 | Flaviviridae |  | Kamiti River virus isolate SR-82 polyprotein precursor, gene, complete cds |
| AJ299445.1 | Flaviviridae |  | Montana myotis leukoencephalitis virus complete genomic RNA |
| AJ242984.1 | Flaviviridae |  | Modoc virus genomic RNA for polyprotein gene |
| AF311056.1 | Flaviviridae |  | Deer tick virus strain ctb30 polyprotein gene, complete cds |
| AF331718.1 | Flaviviridae | Homo sapiens | Alkhurma virus strain 1176 polyprotein gene, complete cds |
| AF326573.1 | Flaviviridae |  | Dengue virus type 4 strain 814669, complete genome |
| AF253419.1 | Flaviviridae |  | Langat virus strain TP21 polyprotein gene, complete cds |
| AF160193.1 | Flaviviridae |  | Apoi virus polyprotein gene, complete cds |
| AF144692.1 | Flaviviridae |  | Rio Bravo virus strain RiMAR polyprotein gene, complete cds |
| AF161266.1 | Flaviviridae |  | Murray Valley encephalitis virus strain MVE-1-51, complete genome |
| L40361.3 | Flaviviridae |  | Tick-borne encephalitis virus-Siberian subtype polyprotein gene, complete cds |
| U87411.1 | Flaviviridae |  | Dengue virus type 2 (strain 16681) polyprotein mRNA, complete cds |
| Y07863.1 | Flaviviridae |  | Louping ill virus, complete genome |
| U88536.1 | Flaviviridae |  | Dengue virus type 1 clone 45AZ5, complete genome |
| U27495.1 | Flaviviridae | Ixodes ricinus | Tick-borne encephalitis virus-European subtype strain Neudoerfl polyprotein gene, complete cds |
| M12294.2 | Flaviviridae |  | West Nile virus RNA, complete genome |
| M91671.1 | Flaviviridae |  | Flavivirus cell fusing agent polyprotein gene, complete cds |
| M18370.1 | Flaviviridae |  | Japanese encephalitis virus (strain JaOArS982), complete genome |
| M93130.1 | Flaviviridae |  | Dengue type 3 virus complete genome RNA, complete cds |
| L06436.1 | Flaviviridae |  | Powassan virus strain LB, complete genome |
| D00246.1 | Flaviviridae |  | Kunjin virus gene for polyprotein (C, prM, E, NS1, NS2A, NS2B, NS3, NS4A, NS4B, NS5), complete cds |
| X03700.1 | Flaviviridae |  | Yellow fever virus complete genome, 17D vaccine strain |
